# Separate functional subnetworks of excitatory neurons show preference to periodic and random sound structures

**DOI:** 10.1101/2021.02.13.431077

**Authors:** Muneshwar Mehra, Adarsh Mukesh, Sharba Bandyopadhyay

**Affiliations:** Advanced Technology Development Centre, IIT Kharagpur; Information Processing Laboratory, Department of Electronics and Electrical Communication Engineering, IIT Kharagpur

## Abstract

Auditory cortex (ACX) neurons are sensitive to spectro-temporal sound patterns and violations in patterns induced by rare stimuli embedded within streams of sounds. We investigate the auditory cortical representation of repeated presentations of sequences of sounds with standard stimuli (common) with an embedded deviant (rare) stimulus in two conditions – Periodic (Fixed deviant position) or Random (Random deviant position), using extracellular single-unit and 2-photon Ca^+2^ imaging recordings in Layer 2/3 neurons of the mouse ACX. In the population average, responses increased over repetitions in the Random-condition and were suppressed or did not change in the Periodic-condition, showing irregularity preference. A subset of neurons also showed the opposite behavior, indicating regularity preference. Pairwise noise correlations were higher in Random-condition over Periodic-condition, suggesting the role of recurrent connections. 2-photon Ca^+2^ imaging of excitatory (EX) and parvalbumin-positive (PV) and somatostatin-positive (SOM) inhibitory neurons, showed different categories of adaptation or change in response over repetitions (categorized by the sign of the slope of change) as observed with single units. However, the examination of functional connectivity between pairs of neurons of different categories showed that EX-PV connections behaved opposite to the EX-EX and EX-SOM pairs that show more functional connections outside category in Random-condition than Periodic-condition. Finally considering Regularity preference, Irregularity preference and no preference categories, showed that EX-EX and EX-SOM connections to be in largely separate functional subnetworks with the different preferences, while EX-PV connections were more spread. Thus separate subnetworks could underly the coding of periodic and random sound sequences.

**Significance Statement:** Studying how the ACX neurons respond to streams of sound sequences help us understand the importance of changes in dynamic acoustic noisy scenes around us. Humans and animals are sensitive to regularity and its violations in sound sequences. Psychophysical tasks in humans show that auditory brain differentially responds to periodic and random structures, independent of the listener’s attentional states. Here we show that mouse ACX L2/3 neurons detect a change and respond differentially to changing patterns over long-time scales. The differential functional connectivity profile obtained in response to two different sound contexts, suggest the stronger role of recurrent connections in the auditory cortical network. Furthermore, the excitatory-inhibitory neuronal interactions can contribute to detecting the changing sound patterns.

## INTRODUCTION

The neural circuitry operates in a manner to maximize the information flow between the environment and the neural circuit, along with decreasing the redundancies in the dynamically varying stimulus (Chechik et al., 2006; Gaucher et al., 2013). To do that, it becomes necessary to understand the statistical structure of the stimuli in order to interact with the environment efficiently (Winkler et al., 2009; Chait, 2020). The intricate statistical features present in the stimuli form a basis for the neural circuitry to identify relevant features, including expected and unexpected rapid changes in the stimuli and enhance their representation up the cortical hierarchy (Aghamolaei et al., 2016; Parras et al., 2017). This assimilated information is further used in the hierarchy possibly to predict future stimuli based on past experiences and interpret meaningful information. (Winkler et al., 2009; Pearce et al., 2010; Chait, 2020).

The ACX neural circuitry uses the varying spectro-temporal statistical features of the stimuli to generate empirical distributions of the multiple components present within the time varying stimulus. Such studies have been performed across multiple modalities like vision (Turk-Browne et al., 2009), motor control (Bestmann et al., 2008), speech (Saffran et al., 1996), and audition (Andreou et al., 2011; Sohoglu and Chait, 2016; Heilbron and Chait, 2018). Within the auditory domain, it is well known that probabilities of stimuli play an important role in response sensitivity (Ulanovsky et al., 2003a; Yaron et al., 2012). Stimuli with dynamic probabilities have extensively been used to study the neural sensitivities across different species and in multiple stations up the auditory hierarchy (Anderson et al., 2009; Malmierca et al., 2009; Taaseh et al., 2011). However, despite a few studies (Yaron et al., 2012; Aghamolaei et al., 2016; Southwell and Chait, 2018), not much is known about the temporal effects over long-time scales especially in the context of repetitions of sound sequences. To maximize the information flow across the hierarchy, neural circuits often show adaptation at a local synaptic level (Yaron et al., 2012; Hershenhoren et al., 2014) both in feedforward and recurrent connections. Through the past studies, it is well known that stimuli that have a low probability of occurrence elicit the highest response, and this has been extensively studied as a deviant selectivity (Khouri and Nelken, 2015), oddball selectivity (Camalier et al., 2019; Mehra et al., 2019; Srivastava and Bandyopadhyay, 2020) and Mismatch Negativity (MMN). A characteristic feature that manifests at the level of a single neuron is a stimulus-specific adaptation (SSA), which is defined as the adaptation in the responses on repeated presentation of the same stimuli. SSA has been studied across all domains, but caveats on the effect of dynamic long time scale temporal changes concerning standards and deviants embedded in the stream of sounds remain to be investigated.

The number of lines of evidence suggests that neurons in the ACX are sensitive to the auditory environment’s statistical regularities. Previous studies have focused on investigating the effect of context on deviant by violating the variety of patterns (Herrmann et al., 2015; Khouri and Nelken, 2015). However, less is known about how auditory circuitry responds to change in patterns over a long time scale in a noisy environment. The current study tries to bridge this gap using standard and deviant tones and noise in Periodic and Random oddball stimuli, trying to shed light on long adaptation time scales and adaptation to entire stimulus structures. We further establish the differences between the neurons’ encoding properties in ACX in response to the different patterned and random structures in sound sequences. A further step in our study aims to determine the role of individual inhibitory interneurons, PV and SOM, on such adaptation. With 2-P imaging, we decipher a possible role of functional connectivity among the excitatory and inhibitory neurons in carrying out the differential encoding of random and periodic sequences at the network level.

## Results

We recorded a single unit, local field potential (n=11 mice, extracellular recordings) and single neuron Ca^+2^ responses (Gcamp6s injected, n=10 mice and Gcamp-Thy1, n=6 mice) separate experiments from the left ACX of the lightly anaesthetized mouse to oddball sound sequences presented to the right ear. The oddball stimuli consisted of a stream of sounds of 15 tokens with 14 standard (S) sound tokens and an embedded deviant (D) sound token. Each sound token was 50 ms long and gaps of 250 ms with the deviant either presented at a fixed position (8^th^ token, Periodic condition) or randomly (2^nd^-15^th^ token, Random condition) in every iteration of presentation of the sequences (Periodic: Fig. 1A, Left and Random: Fig. 1A, Right). We collected responses in pairs with the Standard and Deviant in a set swapped for both the Periodic and Random conditions. One of the Standard and Deviant tokens was either a tone (f, 8485Hz-34kHz) or a broadband noise (*N*, bandwidth 6-48 kHz with equivalent sound level) to mimic a continuous noisy natural environment with important sound occurring with low probability. Recordings were performed to collect responses to 30 iterations of oddball stimuli presented at 3.33 Hz with an inter-iteration gap of 2-8s. In additional 7 mice (extracellular recordings) and 8 mice (Gcamp Thy1 transgenic mice), data of the same type were collected with both S and D as tones (f1 and f2, different frequencies).

**Figure 1.**
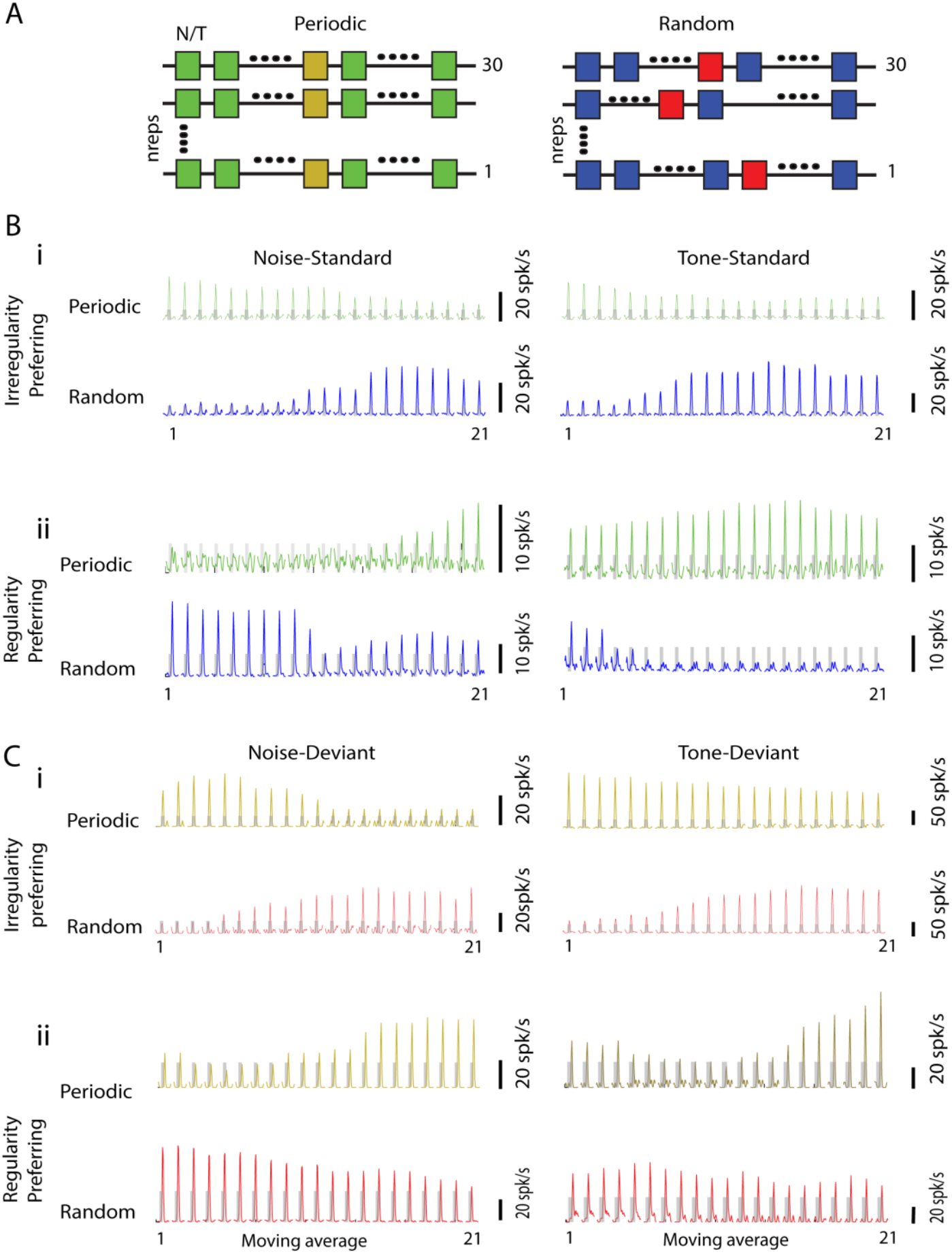
Differential long time scale of adaptation in response to Periodic and Random sound sequences. A) Schematic representation of the Periodic and Random oddball stimulus used in this study. For the Periodic condition, the deviant noise or tone (Yellow, Left) appears once among the standard noise or tone (Green, Left) at 8^th^ position in each iteration with ∼10% probability. In the Random condition, the deviant noise or tone (Red, Right) appears once among the standard noise or tone (Blue, Right) randomly between the 2^nd^-15^th^ position in each iteration. B i) Moving average Psth responses (21 data points obtained from 10 repetitions over 30 repetitions) of a single ACX neuron to the Noise (Left) or tone (Right) as standards in Periodic (Green, Above) or Random (Blue, Below) condition. B ii) Another example showing opposite trend as shown in i), Responses to standard tone and noise in Periodic oddball stimuli increase over 21 moving averages, in contrast to an adaptation in response over later iteration in Random condition. C i) Moving average Psth of a single ACX neuron obtained over 21 moving windows in response to Noise (Left) or Tone (Right) as deviant in Periodic (Above) and Random (Below) conditions. ii) Another example showing the opposite trend as shown in Bii). The colour scheme corresponds to the illustration shown in Fig. 1 A).

### Layer 2/3 single units show that adaptation depends on the structure of a sound sequence

Extracellular single-unit recordings (200-250 µm depth) in the mouse primary auditory cortex show distinct response adaptation patterns to the sound sequences presented in two different contexts of Periodic and Random conditions. We analyzed responses to all the units responding significantly (paired t-test, p<0.05) to at least one of the conditions (Standard and Deviant in Periodic or Random condition). Out of 292 cases obtained for Noise(N)-tone(f) stimuli from 11 mice, 205 cases showed significant responses to standards and deviant in Periodic and Random conditions and were used for further analyses. To observe the nature of adaptation over long time scales, we computed the moving average response in the firing rate observed for Noise (N) and Tone (f) as standard and Deviant, under Periodic and Random conditions (average of 10 successive repetitions moving at step size 1, that is averages of repetitions 1-10, 2-11, …, 21-30, total 21 moving windows, See Methods). For both Periodic and Random conditions, a mean average response to Standard was obtained by considering only the responses to all the standard tokens preceding the Deviant (See Methods). The moving average profile of rate responses of single-units (Fig. 1B-C) and single-channel LFP (Fig. E1-1A-B) response to Standard (Noise; Periodic and Random and Tone; Periodic and Random, Fig. 1Bi-ii) and Deviant (Noise; Periodic and Random and Tone; Periodic and Random, Fig. 1Ci-ii) showed an increase or decrease in response over repetitions. Neurons, like in Fig. 1B-Ci, showing a decrease in average firing rate over repetitions to tone and noise as standard and deviant in Periodic conditions, along with increase or no change in the Random response case are referred to as irregularity preferring. Neurons could also be regularity preferring (Fig. 1B-Cii) when they showed the opposite behavior over repetitions. In the regularity preferring case, the rate response adapted in the random condition over repetitions. On the contrary, response rates increased to tone and noise as standard and deviant in the Periodic condition (Fig. 1B-Cii).

While in the population of single units irregularity and regularity preference (and no preference) was observed, the normalized population moving average response (Figure 2) showed explicit biases. Figure 2Ai shows normalized population mean moving average response (See Methods) to standards and deviant in the Periodic and Random conditions. The population mean moving average response to all standards before the deviant position shows a strong adaptation of responses to tone (Fig. 2Ai, Left, Above) and noise (Fig. 2Ai, Left, below) in the Periodic condition (Fig. 2Ai, Green, Left column). In contrast, in the Random case with an unpredictable deviant position, there is a general increase followed by an adaptation of responses to tone (Fig 2Ai, Left, Above) as well as noise (Fig. 2Ai, Left, Below) as standards (Fig. 2Ai, Blue, Left Column). We also calculated the weighted mean moving average response (See Methods) to tone (Fig. 2Ai, Middle, Above) and Noise (Fig. 2Ai, Middle, Below) to account for bias because of the variable number of standard tokens preceding the deviant in the Random condition (Fig. 2Ai, Middle column, green). The population-weighted mean moving average response shows a similar trend as observed in non-weighted moving average response, Fig. 2Ai (Left Column), where adaptation of responses to tone and noise was observed in Periodic condition contrary to the Random condition. Over repetitions mean moving average response to the Deviant (tone as well as noise) showed lesser or no adaptation compared to the standard in the Periodic condition (Fig. 2Ai, yellow, Right,). Thus, there is likely a higher prevalence of Regularity in Deviant preferring units than in the case of Standards. Similarly, the population mean moving average response to tone (Fig. 2Ai, Right, Above) and noise (Below) as deviant showed a higher increase of firing rate over repetitions in Random case (Fig. 2Ai, Red, Right column) compared to the Periodic condition (Fig. 2Ai, Yellow, Right column). Furthermore, we analyzed RMS values from LFPs (within 50ms sound tokens) corresponding to the channels in which single-units were obtained, to probe for the long time scale of adaptation to tone (Fig. 2Aii, Above) and noise (Fig. 2Aii, Below) as Standard (Fig. 2Aii, Left and Middle column) and Deviant (Fig. 2Aii, Right Column) in Periodic and random cases. We found a similar trend in population mean moving average RMS response to Standards and Deviant in Periodic and random condition (Fig. 2Aii) as observed in single-unit activity (Fig. 2Ai). However, the degree of increase (Periodic_slope_ -Random_slope_) was far higher, as evident, in case of the population single-units instead of LFPs suggesting that such preferences developing over repetitions are emergent in Layer 2/3 (Tone(S): t(204)=2.4065, p=0.0085; Noise (S): t(204)=3.544, p=10e-4; Tone(D): t(204)=2.4216, p=0.0082; Noise (D): t(204)=1.5074, p=0.066, where S: non-weighted normalised Standards).

**Figure 2.**
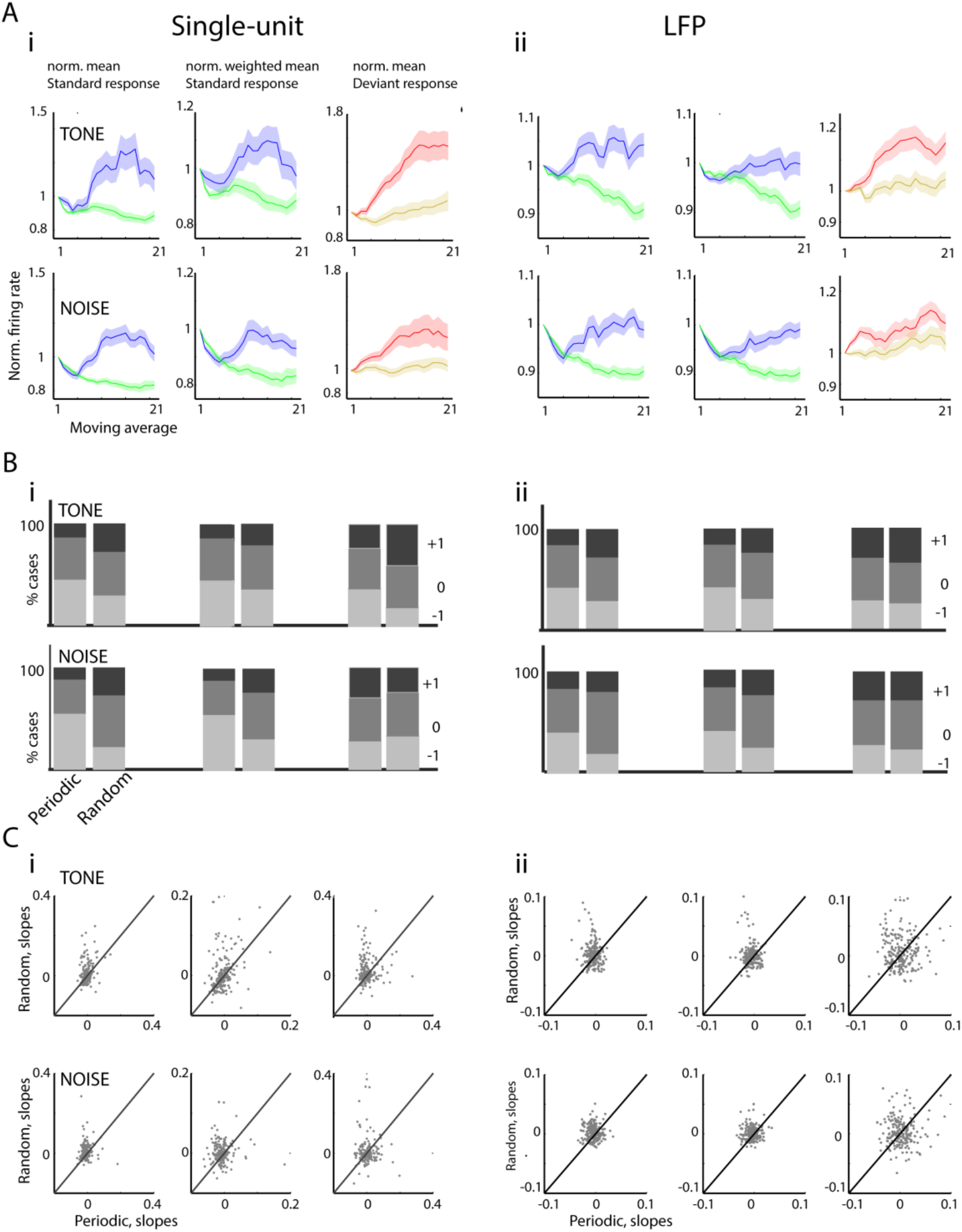
Response adaptation depends upon the structure of a sound sequence in Noise-tone oddball paradigm. A i) (Left) The population mean moving average normalized response rates to Tone (Above) and Noise (Below) as standards in Periodic (Green) and Random (Blue) condition; (Middle) Weighted normalized mean population firing rates to Tone and Noise as standards under Two conditions; (Right) The population moving average normalized response rates to Tone and Noise as deviant in Periodic (Yellow) and Random (Red) condition. A ii) The normalized population mean LFP RMS responses (Left); weighted mean population normalized LFP RMS to tone and noise as standards in Periodic and Random condition (Middle). (Right) Normalized mean population LFP RMS to tone and noise as deviant under two conditions. B i) Proportion of -1, 0, and +1 slopes obtained from normalized and normalized weighted moving average firing rates to tone (Above) and Noise (Below) as standards (Left and Middle) and deviant (Right) in Random and Periodic condition. B ii) Proportion of -1, 0, and +1 slopes obtained from LFP RMS, as shown in B i). C i) The comparison of slopes obtained from the moving average of firing rates in response to tone and noise as standards (Left and Middle) and deviant (Right) under two conditions. C ii) As shown in C i), slopes are obtained from LFP RMS. The slopes obtained to both standard and deviant is significantly higher in Random condition compared to Periodic condition.

To quantify the nature of response change over repetitions, we used linear regression (slopes, see Methods) to account for positive (+1), no change (0) and negative (−1) trends of single-unit activity and LFPs obtained in Periodic and Random case over repetitions (Fig. 2Bi and Fig. 2Bii, respectively). As observed in the mean population moving average response to Standards (Fig. 2Ai-ii, Left and Middle column), we found a higher proportion of neurons showing significant positive slopes in the Random case relative to the Periodic case (Fig. 2Bi, Left and Middle column). Similarly, we found significantly less proportion of neurons showing significant negative slopes for Standards in the Random case compared to the Periodic case (Fig. 2Bi, Left and Middle column. Standard response: Noise, χ^2^=48.44, p<10^−11^; -1(P): p<10^−12^, 0 (R): p<10^−4^, +1(R): p<10^−5^. Tone, χ^2^=16.8, p<10^−4^; -1 (P): p<10^−4^, 0 (R): 0.4, +1 (R): p<10^−4^. Standard weighted response: Noise, χ^2^=25.63, p<10^−11^; -1 (P): p<10^−7^, 0 (R): 0.006, +1 (R): p<10^−4^. Tone, χ^2^=5.47, p=0.06; -1 (P): p=0.03, 0 (R): p=0.36, +1 (R): p=0.02. ‘P’ denotes a higher percentage of the periodic case, ‘R’ denotes a higher percentage in Random cases for corresponding defined slope categories -1, 0, and +1).

In order to define, regularity or irregularity preference, we calculated the change in slope signs (coloured stackbars, Fig. E2-1) by the difference between slopes obtained in the pairs of Random and Periodic conditions (sign(slope_random_)-sign(slope_Periodic_)). Based on the change of slope, we got three categories, regularity preferring (−1, Blue), non-selective (0, Turquoise), and irregularity preferring (+1, Yellow). Thus the relative change in the moving average profile pattern indicated each neuron’s preference to regularity or irregularity over time. A higher percentage of neurons were in the irregularity preferring (Fig. E2-1Ai, Yellow, Left, and Middle bar stacks, Table S2-1) and no-significant change category (Fig. E2-1 Ai, Turquoise, Left and Middle bar stacks) in response to Standards. Between noise and tone, the slope change distribution for noise had more irregularity preferring neurons (Fig. E2-1 Ai, Table E2-1) than tone when presented as Standard, but when presented as Deviant, tone had more irregularity preferring neurons and less regularity preferring neurons (Table S2-1). Thus we found a differential nature of adaptation to tone (Fig 2Bi, Right column, Above) and noise (Fig 2Bi, Right column, Below) as Standard as well as Deviant. When tone was presented as a deviant, the majority of neurons were irregularity selective, as observed in standards (χ^2^=22.26, p<10^−5^ -1(P): p<10^−5^, 0 (R): 0.4, +1 (R): p<10^−5^). On the contrary, when noise was presented as a random deviant, we found fewer neurons with a positive slope than the Periodic case (χ^2^=1.89, p=0.38; -1 (R): p=0.13, 0 (P): p=0.5, +1(P): p=0.12). Similarly, change in slope (sign(slope_random_)-sign(slope_Periodic_)) from Periodic to random also showed a similar differential pattern to deviant tone and noise (Fig E2-1 i-ii, Right column) as observed in the absolute proportions. The above results indicate differential coding of Tone and Noise when presented as deviant under two different contexts.

Similarly, slopes obtained from moving average LFP response to tone and noise as Standard showed significantly higher slopes in Random condition compared to Periodic condition (Fig. 2Bii, Left and Middle, Standard response: Noise, χ^2^=21.95, p<10^−5^; -1 (P): p<10^−6^, 0 (R): p<10^−4^, +1 (R): p=0.22. Tone, χ^2^=12.09, p=0.0024; -1 (P): p=0.0028, 0 (R): p=0.4, +1 (R): p=0.0013. Standard weighted response: Noise, χ^2^=13.47, p=0.0012; -1: p<10^−4^, 0: 0.037, +1: p=0.025. Tone, χ^2^=7.72, p=0.021; -1 (P): p=0.0071, 0 (R): p=0.243, +1 (R): p=0.0166). However, we observed differential activity to tone and noise with higher slopes in response to tone as a Deviant in the random condition in contrast to no-significant slope difference between Random and Periodic condition to noise as deviant (Fig. 2Bii, Right, Noise, χ^2^=1.18, p=0.55; -1 (P): p=-0.15, 0 (R): p=0.18, +1 (P): p=0.5. Tone, χ^2^=1.19, p=0.55; -1 (P): p=0.25, 0 (P): p=0.34, +1 (R): p=0.014). This suggests differential adaptive coding to tone and noise as Deviant when presented in the Periodic and Random conditions. Unlike observed in single unit, the change distribution in LFP showed no significant difference between noise and tone both when presented as standard and deviant (Fig E2-1 Aii, Table E2-1).

A summary of all the recorded single units and recording sites is shown in Fig. 2Ci and Fig. 2Cii, respectively. Fig. 2Ci compares the slopes obtained from moving average single-unit response over repetitions to tone (Above) and Noise (Below) as standards (Fig 2Ci, left and middle column) and deviant (Fig 2Ci, Right) in Periodic (X-axis) and Random (Y-axis) condition. The slopes obtained to standard and deviant in Random condition were significantly higher than slopes in the Periodic condition (Fig 2Ci-ii, Table 1).

**Table 1.** Population summary: Comparison of slope between Periodic and Random sound sequence obtained with Noise(N)-tone(f) oddball paradigm.

|  | Non-weighted Standards (before deviant tokens) |  | Weighted Standards (before deviant tokens) |  | Deviant |  |
| --- | --- | --- | --- | --- | --- | --- |
| Single unit | f | N | f | N | f | N |
| t-test= | 3.8820 | 5.4140 | 2.8425 | 3.6684 | 3.1625 | 1.6678 |
| P val= | <0.0001 | <0.0001 | 0.0024 | <0.0001 | 0.0009 | 0.0484 |
| df= | 204 | 204 | 204 | 204 | 204 | 204 |
| LFP | f | N | f | N | f | N |
| t-test= | 4.1518 | 5.7282 | 3.5667 | 5.4107 | 2.3306 | 0.6627 |
| P val= | <0.0001 | <0.0001 | <0.0001 | <0.0001 | 0.0103 | 0.2541 |
| df= | 204 | 204 | 204 | 204 | 204 | 204 |

So far, we used Noise(N)-Tone(f) oddball stimuli to mimic a noisy ambience with an embedded tone as a Deviant stimulus, as observed in a natural environment (like an adult mouse vocalization in a normal environment). Since deviant selectivity and sensitivity in a sequence of sounds have been previously studied using two tones (f1 and f2), we also used f1-f2 oddball stimulus to investigate if our results hold in such cases as well. We selected f1 and f2 based on frequencies to which significant responses were present. The difference between f1 and f2 varied from 0.25-1.25 octaves. Out of 283 cases obtained from 7 mice, 241 cases showed significant responses to Standards and Deviant in Periodic and Random conditions and further used for analyses. As with the N-f oddball sequences different single units and LFPs in single channels could show irregularity or regularity preference with f1-f2 sequences (Fig. E2-2A). The population mean obtained from single-unit activity followed a similar trend (Fig. E2-2Bi) as observed in N-f oddball stimuli both for Standards and Deviant. The slopes proportion obtained in response to standards and deviant as f1 or f2 in random and Periodic condition followed a similar pattern, as observed in Noise-Tone oddball stimuli when presented as standard (Fig. E2-2Ci; Standard response: χ^2^=23.82, p<10^−6^; -1 (P): p<10^−7^, 0 (R): p=0.002, +1(R): p=0.018. Standard weighted response: χ^2^=22.97, p<10^−7^; -1(P): p<10^−4^, 0 (P): p=0.166, +1(R): p=10^−7^), but not deviant (χ^2^=0.68, p=0.7; -1(P): p=0.33, 0 (R): p=0.2, +1(P): p=0.31). Moreover, the slope comparison between Periodic and random conditions also showed higher positive slopes for standards in random case, and such changes were also observed in slope change (Fig. E2-2Di, Table E2-2).

In the f1-f2 case, the mean population LFP activity varied differentially in Random conditions with a general increase in response to Standards and Deviant, followed by a strong adaptation of responses in later iterations (Fig E2-2Bii). The proportion of slope showed a significant number of +1 slopes for standards in Random condition (Fig. E2-2Cii, Standard response: χ^2^=11.14, p=0.038; -1(P): p=0.002, 0(R): p=037, +1(R): p=0.004), however opposite behavior was observed for Deviant (Deviant response: χ^2^=4.61, p=0.09; -1(P): p=0.36, 0 (R): p=0.035, -1 (−): p=0.032). The comparison drawn between slopes obtained for the Random and Periodic conditions also showed a higher slope for Random conditions standards (Fig. E2-2Dii, Table E2-2). Comparing the normalized population average profiles of LFPs and single units (Periodic_slope_ -random_slope_) again shows that most of the average effect is emergent in Layer 2/3 (f1-f2(S): t(481)=3.371, p=10e-4, p=0.0085; f1-f2(D): t(481)=2.57, p=0.0052) with higher mean absolute slope difference between periodic and random for singe-unit than LFP activity.

In general adaptation in neural responses suggests that the Periodic sequence stream responses should decrease in the successive repetitions. Conversely, if the stimuli structure changes in each iteration, such an adaptation should not occur. Essentially, the present understanding of the adaptation phenomenon supports the existence of the irregularity preferring neurons (higher slope in random, lower slope in periodic case). The primary result obtained with an extracellular single-unit and LFP recordings is that in long time scale adaptation to an entire structure of sound sequence can show decreases or increases in response to the component stimuli over repetitions. We got a sizable number of regularity preferring neurons (Blue, higher slope in Periodic, lower slope in Random, Fig. E2-1; Noise & Tone standard: 24.4%; Noise weighted standard: 22.9%; Tone weighted standard: 23.9%; Noise deviant: 30.24%; Tone deviant: 28.78%; Tone-Tone standard: 23.65%; Tone-Tone weighted standard: 23.03% and Tone-Tone deviant: 28.84%). Such observations indicate that the general understanding of adaptation cannot be easily extrapolated to stimuli consisting of a repeated sequence of sounds, and it warrants further investigation.

We investigated the relation of adaptation to tones in the sequences to the receptive field of neurons. To investigate, we examined the slopes obtained from single-unit moving average responses to tones as Standard and Deviant in N-f (Fig. E2-3Ai-ii) and f1-f2 oddball paradigm (Fig.E 2-3Bi-ii), as a function of distance from the best frequency (BF) of the neuron. In most cases at various distances from the neuron’s BF the mean slopes were either significantly higher or no significant difference (one-sided unpaired t-test) in Random condition compared to the Periodic condition (Fig. E2-3A-Bi-ii). In conclusion, the single-unit and LFP moving average responses to the N-f and f1-f2 oddball stimulus standards show similar tendencies with an increase in response followed by adaptation in Random than in Periodic condition. In contrast, responses to deviant in most cases are affected in a similar fashion, but to a smaller extent.

We also verified slopes difference between Periodic and Random condition by not normalizing with the first moving window (Fig. E2-4A-Ci) and even by considering all the standards (Fig. E2-4A-Cii, Table E2-4) in a given condition. We found our results to be consistent for standards showing higher mean slopes for the Random case. We also see the presence of a substantial number of neurons not showing adaptation to the sound sequences over repetitions in the Periodic case contrary to expectation. Thus we ruled out the effect of normalization and disparity of the number of Standard tokens considered in comparing Periodic and Random sequences. To further evaluate whether cortical structures are necessary to incorporate such long time scales of adaptations, we additionally performed extracellular recordings in the auditory thalamus (n=4 mice, 50 cases). We looked into the slope distributions for noise and tone (Fig. E2-5, Table E2-5a) using Noise-Tone oddball paradigm, as quantified in the ACX moving average data. Unlike the auditory cortex, we did not find a significant difference in Periodic and Random slope distributions in the noise standard stimuli. Still, it was present in tone standard stimuli; these differences were all because of a higher number of negative slopes than the positive slopes, opposite of the auditory cortex. Surprisingly, when the change distribution was considered, we found a higher number of irregularity preferring neurons in noise than tone, when presented as standard, similar to what we saw in the auditory cortex (Table E2-5b).

### Differential Response Strengths of ACX neurons in Random and Periodic Conditions

The ACX neurons are sensitive to statistical structure with higher evoked responses to less probable tones (Ulanovsky et al., 2003b; Malmierca et al., 2009; Yaron et al., 2012). In our study of adaptation to a whole sound sequence structure, we also calculated the CSI index (See Methods) to probe if differences exist between Periodic and Random condition. However, CSI distributions in the population did not show a significant difference in Periodic and Random condition (Kolmogorov Smirnov Test, N-f: p=0.213; f1-f2: p=0.505). A previous study on stimulus structure sensitivity (Yaron et al., 2012) showed higher evoked responses to tones in a Random sequence than Periodic sequences. The observed pattern was dependent on the deviant probability with a stronger effect seen for standards in Random compared to Periodic condition. We examined differences in spike rates in Random and Periodic condition to investigate the spike response dependence at 10% deviant probability in our experimental oddball paradigm.

Contrary to results observed by Yaron et al. 2012, we mostly found higher evoked responses to standards and Deviant in Periodic case relative to the Random condition. Figure 3A summarizes the sound-evoked spike activity to N-f oddball stimuli. The coloured filled circles represent cases where a significant difference (paired t-test, p<0.05) of sound-evoked activity was observed between Random and Periodic condition.

**Fig 3.**
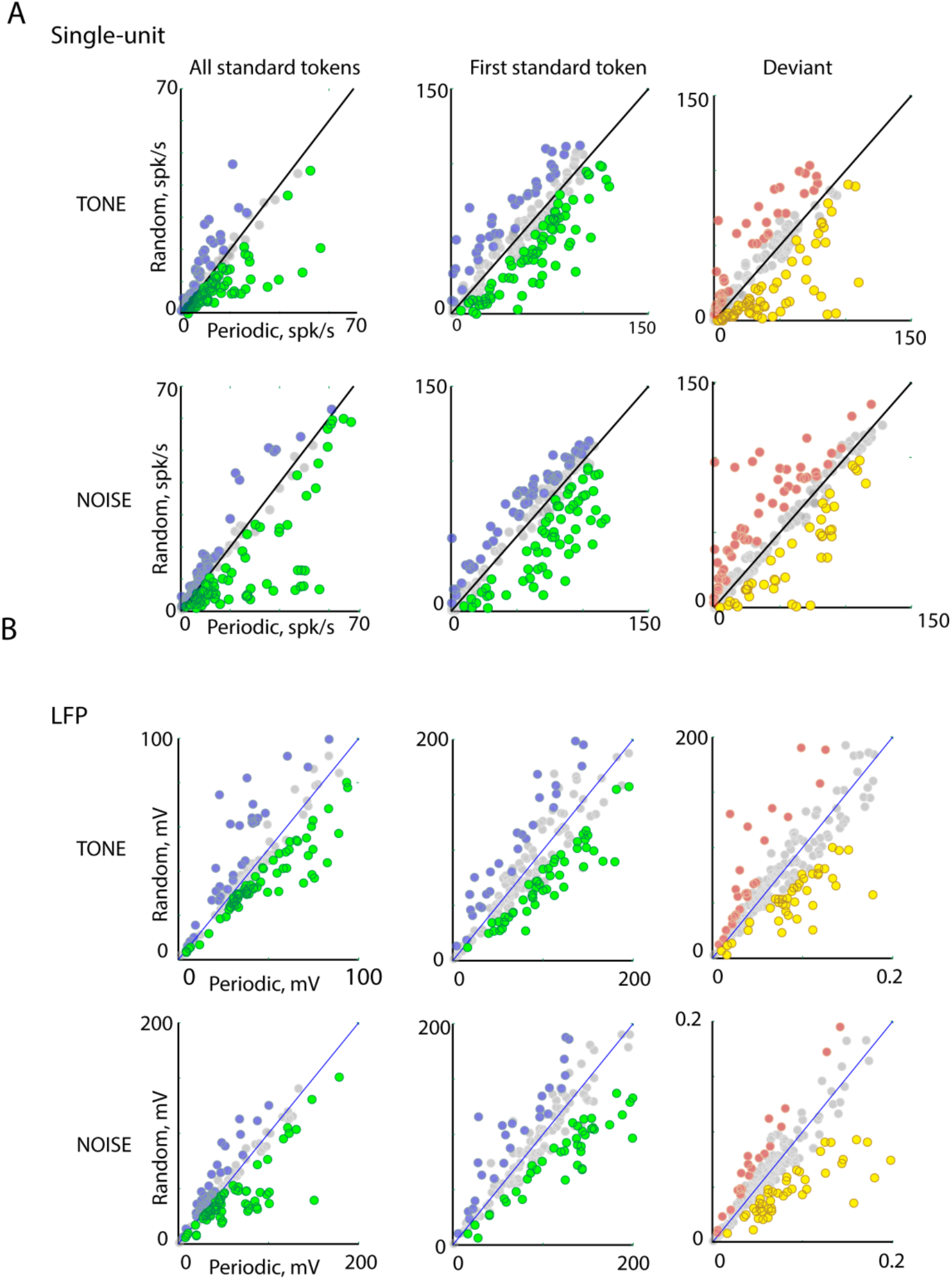
ACX L2/3 Single-unit and LFP responses to the periodic and random sequence. A) Top two rows: Population mean firing rate of cases to all Tone (Above) and Noise (Below) standard tokens in Periodic and random condition (Left). Population mean firing rate of cases to first Tone (Above) and Noise (Below) standard token in Periodic and random condition (Middle). Population mean firing rate to Tone and Noise as deviant under two conditions (Right). B) Bottom two rows: Population mean RMS (LFP activity within stimulus duration) to tone and noise as standard (Left and Middle) and deviant (Right) in Periodic and Random condition. The coloured filled circles represent cases where a significant difference (paired t-test, p<0.05) of sound-evoked activity was observed between Random and Periodic condition.

We calculated spike rates for standards under two conditions. Firstly, by considering a mean response to all the standard tokens in a presented oddball sequence (Fig. 3A, Left) and secondly, by considering only the mean response to the first token (Fig. 3A, Middle), as robust responses were observed at first token followed by strong adaptation to successive tokens. Considering all the standard tokens, the responses to both Standard tones (t(204)=3.35, p<0.0001) and Standard noise (t(204)=4.34, p<0.0001) in the Periodic condition were larger compared to the Random condition (Fig. 3A, Left, Table 2). We found a higher proportion of cases showing higher evoked responses to tones (146/205, 71%) relative to noise (115/205, 56%) in Periodic condition relative to Random condition. However, the differential response profile was observed with higher rates observed to Deviant tones in Periodic condition (t(204)=2.14, p=0.016), while we saw no significant change to noise as Deviant in Periodic or Random condition (t(204)= -0.79, p=0.786). Whereas, in the LFP recordings, the responses to both Standard tones and noise were mostly larger in the Periodic condition (Left, Standard Tone: t(204)=1.653, p=0.049; Standard Noise: t(204)=1.862, p=0.032, Fig. 3B, Bottom two rows). Similarly, Deviant tone and noise in Periodic condition (Right, Deviant Tone: t(204)=1.658, p=0.041; Deviant Noise: t(204)=1.862, p<0.0001, Table 2, Fig 3B) evoked higher potential changes compared to Random condition.

**Table 2.** Population Summary: Responses to Tone and Noise as standard and deviant in Periodic and Random sound sequences

|  | Standards (all tokens) |  | Standards (first token) |  | Deviant |  |
| --- | --- | --- | --- | --- | --- | --- |
| Single unit | f | N | f | N | f | N |
| Number of cases<br>Periodic>Random | 146/205 | 115/205 | 125/205 | 96/205 | 107/205 | 91/205 |
| Fraction | 71% | 56% | 61% | 47% | 52% | 44% |
| t statistic | 3.35071 | 4.3363 | 3.1197 | 2.8914 | 2.1480 | -0.7960 |
| df | 204 | 204 | 204 | 204 | 204 | 204 |
| P val | <0.0001 | <0.0001 | 0.0010 | 0.0021 | 0.0164 | 0.7865 |
| LFP |  |  |  |  |  |  |
| Number of cases<br>Periodic>Random | 141/205 | 115/205 | 132/205 | 123/205 | 109/205 | 124/205 |
| Fraction | 69% | 56% | 65% | 60% | 53% | 61% |
| t statistic | 1.6537 | 1.8624 | 2.3701 | 4.1438 | 1.6580 | 3.3100 |
| df | 204 | 204 | 204 | 204 | 204 | 204 |
| P val | 0.0499 | 0.0320 | 0.0094 | <0.0001 | 0.0413 | <0.0001 |

Likewise, spike rates obtained using tone (f1)-tone (f2) stimuli in Periodic and Random condition yielded a similar response profile as observed in Noise-tone stimuli. We found similar results for the single-unit activity for Standards (Fig. E3-1A, Left-Middle) as well as Deviant (Fig. E3-1 A, Right). A majority of cases with a significant difference between the two conditions responses showed higher responses in the Random condition (standard: 304/482, 64%, deviant: 280/482, 58%). Also, overall rates were found to be higher in Periodic condition to both Standards (t(481)=4.69, p<0.0001) as well as deviant (t(481)=2.39, p=0.008). Moreover, consistent results were obtained with LFP activity, showing higher evoked response to standards (t(481)=3.37, p=0.0003) and deviant (t(481)=5.58, p<0.0001) tones in the Periodic condition (Fig E3-1. B, Table. E3-1). Overall, the firing rate and input synaptic activity reflected by LFP suggests that the ACX neurons are sensitive to structures within the Random and Periodic sequences and induces higher average evoked responses to the predictable environment than to the Random context in our investigated oddball paradigm. Thus, in the population while slopes indicated a general tendency of responses to rise over repetitions in the Random condition compared to the Periodic condition, the average rates were lower in the Random condition than the Periodic condition suggesting a form of gain control. In order to accommodate a rise in the response rates over repetitions for the Random case, a lowering in the overall rates compared to the Periodic case is achieved. It may be observed in the population average normalized rate response moving average profiles (Fig. 2) that the mean responses are some in both cases in the starting few moving averages and then the responses in the two conditions diverge. Such a mechanism is likely achieved through the network with recurrence (Yarden and Nelken, 2017) or through inhibitory neurons adaptation (Natan et al., 2015a) or gain control (Seybold et al., 2015), or both. We next consider both these factors in extracellular single-unit responses and imaging based single neuron Ca^2+^ responses.

### Random and Periodic condition differentially modulates noise correlations and functional network connectivity

The evoked noise correlations, that is, a trial-by-trial variability between simultaneously recorded neurons decreases after stimulus presentation (Oram, 2011), adaptation (Gutnisky and Dragoi, 2008), and different behavioral and attentional states (Zhang et al., 2014; Francis et al., 2018). Cortical neurons improve the efficiency of encoding of stimulus by minimizing noise correlations (Averbeck et al., 2006; Winkowski et al., 2013). Here to consider the effect of functional recurrence in the network, we investigate the influence of stimulus structure on noise correlations. We first computed the noise correlations between simultaneously recorded pairs of single units in response to the Periodic and Random stimuli. The distance between the pairs of neurons was computed based on the electrodes relative locations in the array from the respective isolated single units. A linear fit on the noise correlation values with distance for both these stimuli showed an expected decrease in noise correlations as a function of distance (Fig 4ABi) observed in the ACX (Rothschild et al., 2010; Winkowski et al., 2013). The mean noise correlation decreased with distance both for the noise Standard (Periodic: p<10e-8; Random: p<10e-12, Fig. 4Ai) and tone Standard (Periodic: p<10e-10; Random: p=0.017, Fig. 4Bi). We next compared the slope between the Periodic and the Random cases and we found that in the case of noise Standard, the drop in the value of noise correlation with increasing distance was sharper for the Random stimuli than the Periodic stimuli (more negative slope for Random; slope comparison t-test t(1394)=2.0567, p=0.019, Fig. 4Ai). However, there was no difference in the case of tone Standard (slope comparison t-test t(1394)=1.187, p=0.117, Fig. 4Bi).

**Figure 4.**
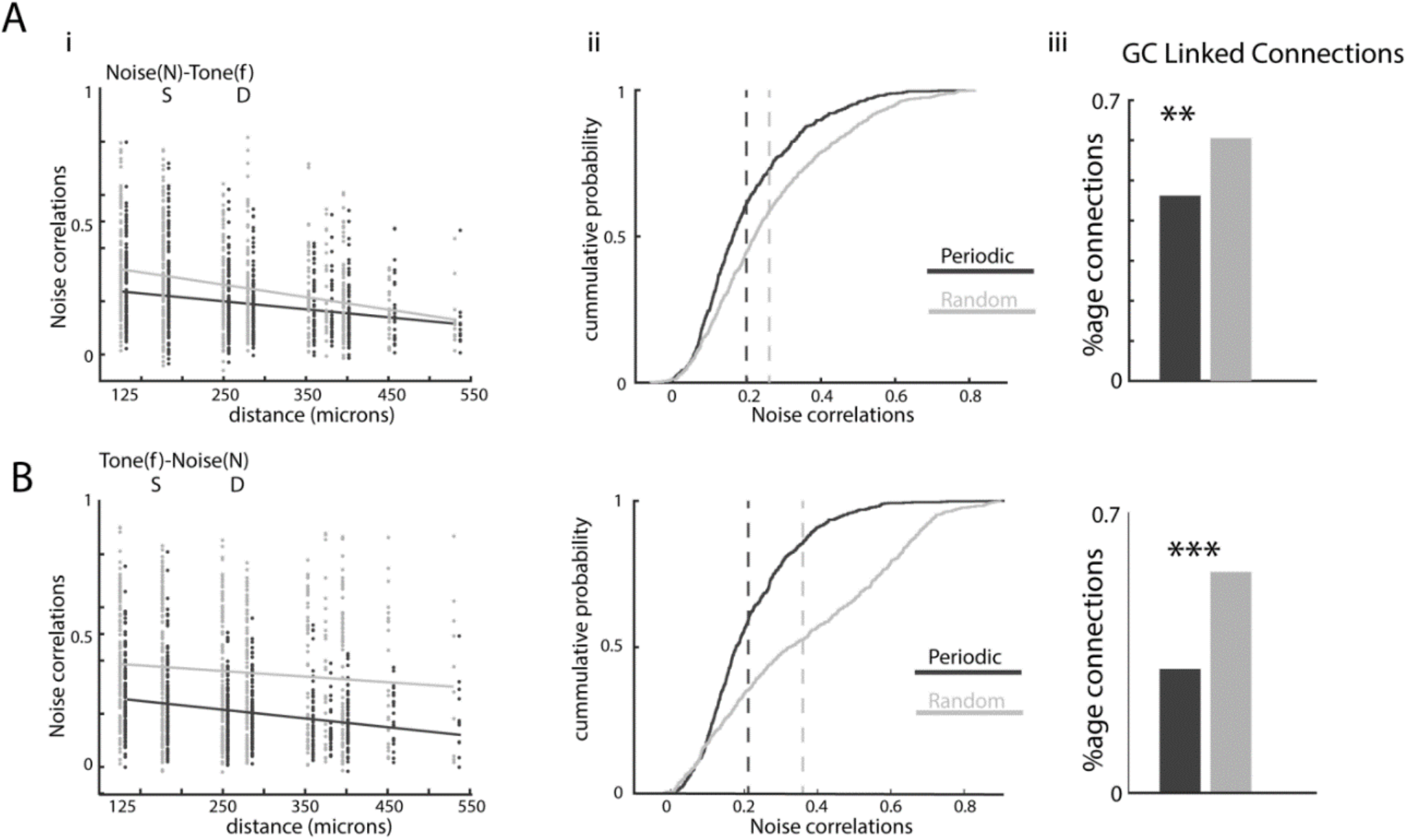
Random and Periodic condition differentially modulates Noise correlation and network connectivity. A i) The noise correlation as a function of the distance between simultaneous recording ACX L2/3 neurons. The gray and black dot indicates the noise correlation obtained between simultaneously recorded neurons in response to Noise (Standard)-tone (Deviant) stimuli in Random and Periodic condition, respectively. The gray and black line shows the linear regression on the distance for both the stimulus in Random and Periodic conditions (Left). ii) Cumulative distributions of Noise correlations for Random and Periodic condition show higher noise correlation in Random deviant context compared to Periodic condition (Middle). iii) Connection probability based on Granger Causality in Periodic and Random condition. B) Same as shown in Ai-ii, Noise correlation and proportion of connected pairs obtained between a pair of simultaneously recorded neurons in response to Tone (Standard)-Noise (Deviant) stimuli in Random (gray) and Periodic (Black) condition. iii) Connection probability based on Granger Causality in Periodic and Random condition. (z-proportionality test, **p<0.01 and ***p<0.001)

We next compared the noise correlations in pairwise responses of single units between the Random condition and the Periodic condition for both kinds of stimuli, Noise(Standard)-Tone(Deviant) and Tone(Standard)-Noise(Deviant) (Fig. 4A-B, Above and Below, respectively). We found the mean of the noise correlations under Random conditions were higher than that under the Periodic condition for both in the noise standard case (paired one-sided t-test t(698)=9, p<10e-17, Fig. 4Aii) and tone standard case (paired one-sided t-test t(698)=19.54, p<10e-67, Fig. 4Bii). Granger causality analysis also revealed a higher proportion of connections involved in Random condition compared to Periodic condition (Fig. 4 A-Biii, z –proportionality test).

Likewise for tone(f1/f2, S)-tone(f1/f2, D) stimuli, mean noise correlation for the Random stimuli was higher than that of the Periodic stimuli (paired one-sided t-test t(1397)=31.925, p<10e-167, Fig. E4-1 ii). In a comparison of the slopes, both for the Periodic and the random stimuli, the noise correlation decreased with increasing distance (Periodic: p<10e-3; random: p<10e-4), but there was no significant difference between the slopes (slope comparison t-test t(2792)=0.058, p=0.476, Fig E4-1i). Our results indicate that the structure of stimuli modulates noise correlations by showing higher noise correlation in random sequences compared to the Periodic stimulus. It essentially indicates that functional connectivity increases during the randomness in structure. Thus pairwise functional connections loosely inferred from noise correlations suggests sparse connectivity in the network during presentations of a periodic stimulus compared to its random counterpart.

### 2-photon Ca^+2^ imaging shows the presence of regularity and irregularity preferring Excitatory as well as Inhibitory neurons

The scaling up or down of the neuronal rates in response to Periodic and Random stimuli hints towards a control mechanism to regulate the neural circuit’s information flow. The role of inhibitory neurons has been implicated in the suppression of neuronal activity (Kato et al., 2017), gain control (Ferguson and Cardin, 2020), and correlated neural fluctuations (Okun and Lampl, 2008). Given all our observations so far, we examined the role of the two most studied interneuron classes in the ACX, namely PV and SOM. We performed in-vivo 2-photon Ca^+2^ imaging in Layer 2/3 of the Gcamp-6s injected (PV-Cre [JAX 008069], SOM-IRES-Cre [JAX 90 013044], ROSA LSL-tdTomato [JAX 007908) mouse (EX: Non-PV, 4mice and Non-SOM, 6 mice; 23 ROI’s, Fig. 5A) auditory cortex in response to Periodic and Random stimuli. For our analyses of imaging data, we considered mean df/f only for Standards preceding deviants for response mean. Due to scan rate constraints, we could not obtain the response to the Deviant embedded in the stream reliably. Since only responses to Standards are considered in the imaging results, we refer to noise Standard and tone Standard responses as simply ‘Noise’ and ‘Tone’, respectively. All neuronal populations, EX, PV and SOM, showed the different categories of effects on response adaptation over repetitions of the two types of sound sequences, as observed with single units (Fig. 5B, Noise(Above), Tone (Below)).

**Figure 5.**
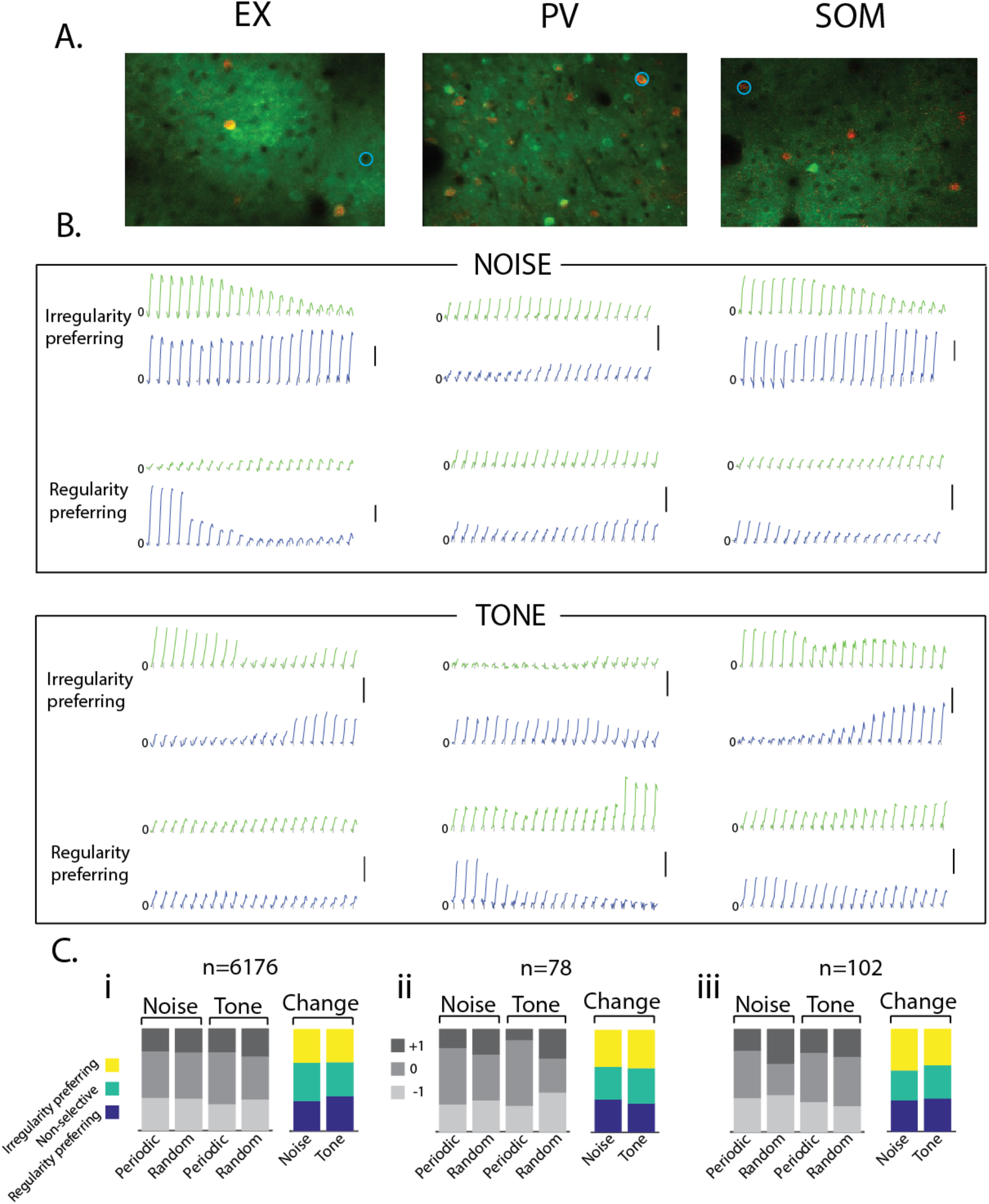
2-P Ca^+2^ imaging response for Periodic and Random stimuli in Excitatory and Inhibitory neurons. A. 2-P Ca^+2^ images showing three types of interneurons, namely, Excitatory (EX), Parvalbumin (PV) and Somatostatin (SOM). Respective neurons are marked in circles. B. Example traces of the three populations with two different types of behavior each (regularity preferring and irregularity preferring) shown in a moving average fashion with a window of 10 frames and an increment of 1 frame (making a total of 21 windows) The black vertical line represents 20% change in df/f. C. Slope distributions (gray stack bars) for the two types of stimuli (noise and tone) when presented in periodic and random fashion and the change distributions (colored stack bars) showing the proportions of regularity preferring, non-selective and irregularity preferring neurons for i) EX, ii) PV and iii) SOM.

We separated the responses into three categories, based on the sign of the slope of the across iteration moving averaged response for both Periodic and Random conditions, positive (+1), no change (0), and negative (−1). Within the EX population, we found a significant difference between the slope distribution in response to Periodic versus Random stimuli for tone Standards (χ^2^ multi-proportion test, χ^2^=97.36, p<<10e-16), but not for noise Standards (χ^2^=1.75, p=0.4). Furthermore, this change in the distribution was due to a higher percentage of both positive and negative slopes and a lower percentage of no-change slopes in the Random case as compared to the Periodic case (one-sided z-proportionality test, -1: p<10e-9; 0: p<10e-16; +1: p<10e-8, Fig.5Ci, gray bars). A similar trend was found in PV neurons both in tone Standards (χ^2^=16.11, p<10e-4) and noise Standards (χ^2^=1.77, p=0.41). In PV neurons, the change in tone standard slope distribution had a contribution of a higher number of both positive and negative slopes and a lower number of neutral slopes (one-sided z-proportionality test, -1: p=0.04; 0: p<10e-5; +1: p=0.002, Fig. 5Cii, grayscale bars). Contrary to EX and PV neurons behaviour, the SOM neurons registered a change in slope distributions in case of noise Standard (χ^2^=6.48, p=0.04) and not in tone Standard (χ^2^=0.6, p=0.73). An increase in the negative number slopes and a decrease in the number of neutral slopes drove this change, while the proportion of positive slope roughly remained the same (−1: p=0.33; 0: p=0.01; +1: p=0.02), (Fig, 5Ciii, grayscale bars).

As we did with the extracellular recordings, we further obtained distributions of the difference in sign of slopes of across iteration moving averaged response for Periodic vs Random conditions (sign(slope_Random_)-sign(slope_Periodic_)), and got the same three categories, Regularity preferring, Non-selective and Irregularity preferring. Within the excitatory population, the difference distribution for noise standard was significantly different from tone standard (χ^2^=38.1, p<10e-8). It was because of more number of regularity preferring and less number of non-selective neurons in the standard tone distribution as compared to the standard noise distribution. The number of irregularity preferring neurons remained the same both in noise standard and tone standard (Regularity preferring(tone): p<10e-8; Non-Selective (noise): p<10e-6; Irregularity preferring (tone): p=0.3), (Fig. 5Ci, colored bars). In the PV population, the difference-distributions of noise standard and tone standard roughly remained the same (χ^2^=0.286, p=0.866),(Fig. 5Cii, colored bars), and so was with SOM neurons (χ^2^=0.53, p=0.76), (Fig. 5Ciii, colored bars).

To rule out the possibility of the results being influenced by the Gcamp-6s longer Ca^+2^ dynamics used above, we performed additional experiments on Gcamp-Thy-1 (EX, n=6 mice, 10 ROIs) transgenic animals (Fig. E6-1, A). Similar to the Gcamp-6s injected animal response profiles, the slope distribution for tone were different for periodic and random case (X^2^=71.4, p<10^−16^), with more number of positive slopes and less number of negative slopes (−1: p<10e-13; 0: p0.2; +1: p<10e-12) in Random case. However, unlike GCamp-6s injected animal slope response profiles, we got the significant difference between periodic and random case for the noise stimuli also (χ^2^=52.27, p<10^−12^), with more number of negative slopes and fewer positive slopes in the random case (−1: p<10e-12; 0: p=0.04; +1: p<10e-7) (Fig. E6-1 Ai). We got the same result when we presented the tone(f1)-tone(f2) stimuli (n=8, Gcamp Thy1 transgenic mice, 7 ROIs, Fig. E6-1), where the distribution of slopes was significantly different (χ^2^=39.12, p<10^−9^). This was fueled by more number of positive slopes and fewer negative slopes in the random case than the Periodic case (−1: p<10^−8^; 0: p=0.127; +1: p<10^−6^), (Fig. E6-1 Aii). We found consistent observation of higher positive slopes for tone Standards in Random condition in injected Gcamp6s and Gcamp-Thy1 (Noise-tone and tone-tone stimuli presented) transgenic animals and also the presence of a sizeable number of Regularity preferring neurons (Fig. E6-1B).

We compared the mean df/f values between Periodic and Random stimuli for the three neuron types in response to both noise and tone. Unlike in the single unit mean rate responses (Fig. 3), the mean df/f for the Periodic case was the same as that of the Random case, except the PV neurons for noise which showed a significant difference (Fig. E5-1, Table E5-1). This could be due to the difference in rate and mean df/f not being identical due to the effects of saturation or biased sampling with electrodes compared to 2-photon imaging. We also see that SOM neurons behavior within its respective population is different from that of PV and EX. The difference in the distribution of slopes for Periodic and Random was significant for tone standard in PV and EX. However, we observed significant slope difference between the two conditions for Noise standards in SOM neurons. These differences could arise because of the differential connectivity shown by PV and SOM neurons across columns and their bandwidths (Kato et al., 2017). Thus specific neuronal types (EX or PV-SOM) did not show the prevalence of a particular type of adaptation quantified through the slope of response change over repetitions. Also, we observed a dramatic shift in noise correlations under the two conditions (Fig. 4B), with higher noise correlations in nearby neurons (Fig. 4A). Thus we hypothesize that the local functional recurrent connectivity could underlie the different slope categories or slope change categories – regularity, irregularity preferring or non-selective. Hence, we next investigated the nature of adaptational changes over repetitions in the Random and Periodic slope distributions with respect to the recurrent connectivity within the micro-circuitry.

### Differential connectivity profile of PV and SOM in response to Periodic and Random oddball stimuli

Next, we looked into the connectivity profiles within various subpopulations of neurons imaged. We classified each pair of simultaneously imaged neurons into two categories, either connected or not connected. The classification is based on comparison of bootstrapped noise-correlations between pairs of cells and chance noise correlations estimated from the same paired data by destroying the simultaneity of the responses (order randomized) in the pairs of neurons (See Methods). Followed by this, we divided the connections into five different classes based on the neuron types in the pair, namely EX-EX, EX-PV, EX-SOM, PV-PV, and SOM-SOM. Since each cell could respond differently to the noise Standard and tone Standard stimuli, we categorized these cells based on the sign of the slope of the across iteration moving averaged response. Thus, we got 9 categories (3 slope types for noise standard multiplied by 3 slope types of tone standard), for each of the two conditions -Periodic and Random. We constructed a weighted graph consisting of 9 nodes for the 9 categories, and the edge weights denoting the total number of connections between them as estimated above. We used Gini-index (Goswami et al., 2018) as a measure of sparsity (See methods) to quantify this connectivity between the 9 different categories (a higher Gini-index signifies higher sparsity in the connectivity, which implies weaker and lesser connectivity between neurons of different categories). We bootstrapped the connectivity graphs to generate several replicates (500) and constructed 90% confidence intervals for the Gini-index. In the case of EX-EX connectivity, the sparsity of connections in the Periodic condition was significantly higher than that in the Random condition (Fig. 6Ai) implying higher functional connectivity among the different slope category neurons in case of Random stimuli. We found a similar pattern in the case of EX-SOM (Fig. 6Aii, Right) connections but was opposite in EX-PV connections (Fig. 6Aii, Left). The PV-PV and the SOM-SOM connections did not show significant differences between the Periodic and random cases within the inhibitory subpopulation (Fig. 6Aiii). To obtain the Gini-index over repetitions to see how the functional network connectivity changed, we computed it in moving windows of 10 successive repetitions (Fig. 6B). We see that the observed difference in Gini-indices between the Periodic and Random conditions was maintained within the EX-EX population from the initial 10 iterations to the final 10 iterations (Fig. 6Bi). Similar results were found in EX-EX connections (Thy1 positive EX neurons) from the data obtained in GCamp-Thy1 transgenic mice with both the Noise-Tone stimuli (Fig. E6-1Ci) and the Tone-Tone stimuli (Fig. E6-1Cii). However, the above was not the case with the EX-PV and EX-SOM connections, where we found that the difference was absent in the initial 10 iterations, but emerged later and was evident in the last 10 trials (Fig. 6Bii). Thus the change in functional recurrence in the inhibitory-excitatory connections emerged over the sequence repetitions with EX-PV connections showing a different connectivity nature under the two conditions. The PV-PV and SOM-SOM function connectivity remained the same throughout the repetitions (Fig. 6Biii). The above results indicate that the two inhibitory neuronal types change their effective influence on the network differently under the two stimulus conditions with EX-PV connectivity becoming largely sparse under the Periodic condition compared to the Random condition and that such effect emerges over time with increasing repetitions. The results of Gini-index also illustrate that functional connections exist within similar slope categories in EX-EX and EX-SOM but not EX-PV. PV-PV and SOM-SOM connections are not discussed further as firstly there was no differential effect observed in them for the two stimulus conditions. Secondly, our dataset does not have simultaneous recordings of large numbers of PV or SOM neurons, with few (2-5) PV or SOM neurons in every ROI investigated.

**Figure 6.**
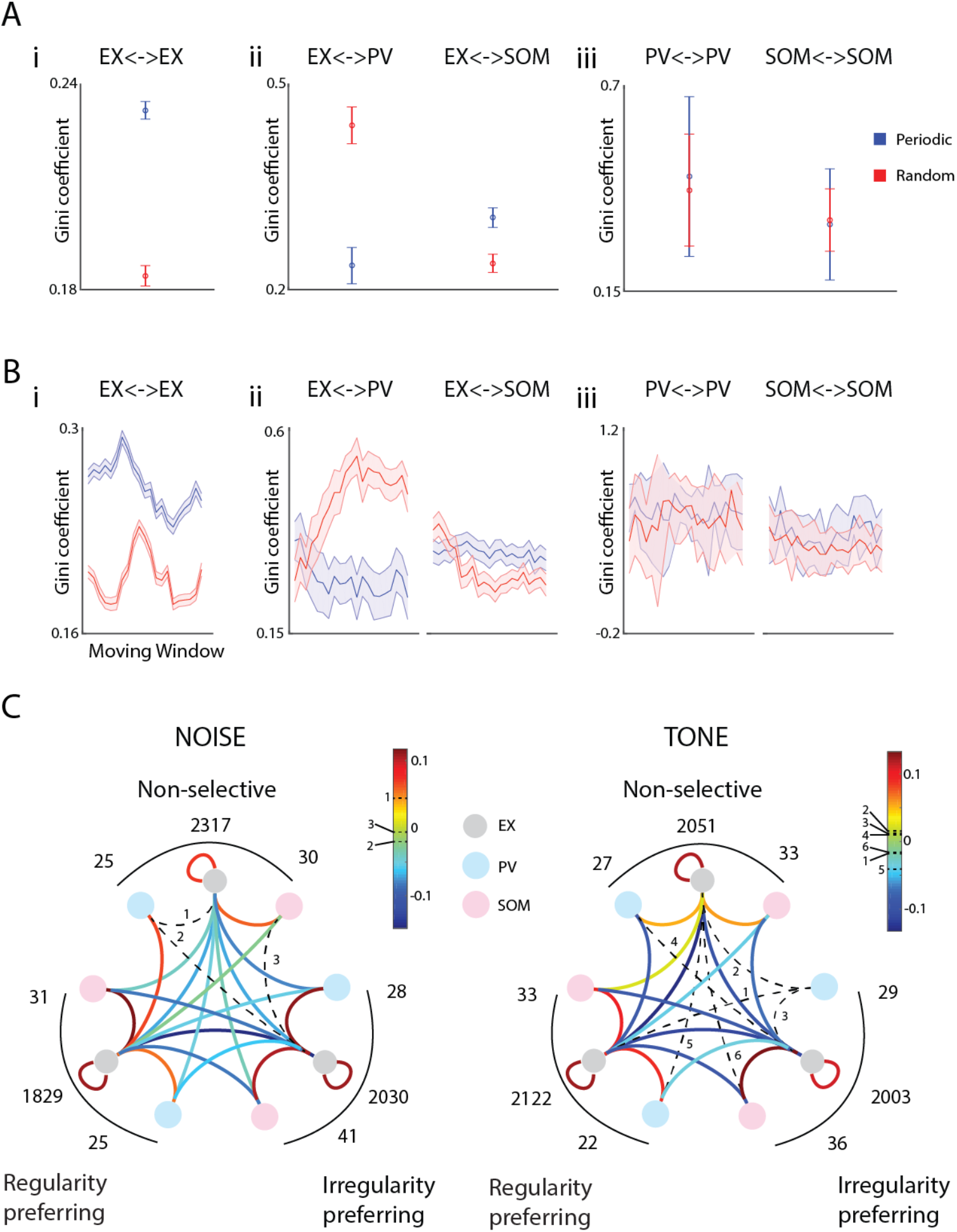
Differential connectivity profile of PV and SOM in response to Periodic and Random oddball stimuli. A. Mean Gini coefficient calculated for ajj the 30 stimuli iterations. i) within the ex-ex population. ii) within the EX-PV and ex-som population. iii) within PV-PV and som-som population. B. Gini coefficient calculated across the iteration in a moving average window of 10 iterations, with the shaded bar representing the 90% confidence interval. i) within the ex-ex population. ii) within the EX-PV and ex-som population. iii) within PV-PV and som-som population.. C. Network schematic showing the relative probability of connectivity with respect to chance between the two nodes representing the two cells with their respective slope change behavior for both noise and tone stimuli. The solid-coloured lines represent the significant connections (see methods), with the color representing the relative probability value. The dotted lines represent connections with the non-significant value of relative probability, with the numbers representing their respective values on the color bar. The numbers along the arc represent the number of cells in the respective category.

To further elucidate the nature of the micro-network in terms of preference of neuron types to regularity and irregularity we quantified the connections between the excitatory and inhibitory neurons (EX-EX, EX-PV and EX-SOM) of various slope change categories. Thus we constructed 99% confidence intervals of connection probability between two given slope change categories and compared it with the observing a connection by chance in the same pairs of neurons by randomizing the connectivity between pairs (See methods). We obtained the relative probability of occurrence of connections between pairs of neurons (type and category) with respect to chance for all the possible connection categories (6 for EX-EX pairs and 9 for EX-PV and EX-SOM pairs) and if they were significantly higher (warm color edges, Fig. 6C) edges or lower (cold color edges, Fig. 6C) than chance or were at chance level (dashed edges, Fig. 6C). We found that the connections within the same slope change categories were much more probable within the EX-EX pairs than across slope change categories for both the noise standard (Fig. 6C, Left) and tone standard stimuli (Fig.6C, Right). A similar trend was found in the EX-SOM connections also, while the relative connection probabilities in the EX-PV population were more spread than the former two populations both within and across categories, with many non-significant connections in the tone standard stimuli. It is evident that significantly higher than chance connectivity was mostly present within a category (warm color edges within outer arcs representing preference categories), while across category connections were mostly significantly lower than chance (cold colored edges make outside category connections). Departure from the above was mostly due to the involvement of PV neurons. The summary of connections between slope change or regularity/irregularity preference categories (Fig. 6C) shows that separate functional subnetworks of EX-EX and EX-SOM are present that show regularity or irregularity preference. These average networks are functionally modulated by PV neurons primarily from outside category and SOM from within category over repetitions differentially under the two stimulus conditions (Fig. 6AB).

## DISCUSSION

Our results show that the auditory cortical system, both at the individual neuron level and local circuit level, is susceptible to the sensory stream’s statistical structures. The degree of randomness in the deviant positions in the two stimuli we used had far-reaching consequences, which differentially influenced the neurons’ responses. Such differences may shed light on the various ways through which the brain represents structures to enhance discriminability among the various components of the auditory stimuli.

Stimulus-specific adaptation corresponds to the local short-term adaptation effect on the repeated presentation of a periodic stimulus (Ulanovsky et al., 2003a; Pérez-González and Malmierca, 2014). We also found a long-scale version of adaptation resulting from a repeated presentation of a similar stream of stimulus tokens where we consider adaptation to the entire sound structure. These structural differences pertaining to the position of the deviant token between the successive streams essentially shaped the degree and nature of adaptation occurring in the neural circuit, and the results likely indicate that previous history of the stimuli is stored at multiple timescales and is used by the neural circuit to shape its future responses.

Previous studies used a continuous stream of stimuli tokens, which allowed both the short term (because of successive tokens) and long term (because of the probabilistic structure) synaptic depression to co-exist (Ulanovsky et al., 2003a; Yaron et al., 2012; Malmierca, 2014). We presented the oddball stimuli in a successive iteration fashion with sufficient gap (2-8s) between neurons, thereby allowing them to recover from the local short-term synaptic depression and form a separate long-stream memory of the deviant position. We seek to understand if auditory circuitry keeps track of the previous history of structure in a stream of sounds. In an attempt to investigate changes over the long-timescale, we probed change in firing rates and synaptic input activity reflected by LFP across multiple iterations to a Periodic and random stream of sounds in L2/3 neurons of the auditory cortex (Fig. 2A-C). We found a higher number of positive slopes for responses to standards and deviant tones obtained over iterations presented in the Random condition compared to the Periodic condition (Fig 2B-C). Such results imply that the degree of adaptation for successive repetition of the complex randomly-patterned stream of sounds was much higher as compared to the successive repetition of the Fixed-patterned periodic stimulus. Furthermore, standard responses were more affected than the deviant responses, which reflects context-specific modulation. Taken together, these changes reflect long-term memory mechanism; where ACX neuron continually tracks and compares the incoming sensory patterns to past history and accordingly update the future response.

We further found rates for the standards and deviant in Periodic conditions to be higher compared to a random deviant case (Fig 3A-B). In our study, the rate-specific results observed under two conditions were contrary to Yaron et al., 2012 study, which could be due to the long-stream stimulus presented in earlier study contrast to oddball stimuli used in our study with sufficient gap between each iteration. However, our results are in accordance with higher evoked potential for the outliers in regularly-patterned (REG) stimulus compared to Random (RAND) tone pip sequences observed in a study by Southwell and Chait, 2018. Our pieces of evidence further support results obtained in the musical expectation studies (Vaz Pato et al., 2002; Pearce et al., 2010), where they observed a higher deviant response within the structural context compared to Random context. Since experiments were performed under lightly anaesthetized condition, it is unlikely to consider the involvement of attention (Zhao et al., 2013) in such experimental paradigms. Thus, our results for long time scale adaptation changes and differential rates to Periodic and Random condition suggest long-term structure learning in auditory cortex neurons based on the past experience, as a new structure emerges in every iteration in the Random condition.

We observed a higher degree of functional connectivity in the random deviant stimuli compared to the fixed deviant (Fig. 4A). Even though noise correlation gives us a relative measure of the connection strength, it can be conducive to substantiate our results comparing the effect of stimuli complexity on the local circuits interconnectedness(Ginzburg and Sompolinsky, 1994; Averbeck et al., 2006). We found a higher value of mean noise correlation in the random deviant stimuli compared to a periodic sequence. Although expected, it gives us a hint that complex stimuli structures recruit a larger number of neurons within different levels of hierarchy to create a “memory” and optimize their processing to show the respective adaptations. This result holds for both the noise-tone and tone-tone stimuli, essentially implying that the nature of stimulus has little to do with the density of active circuit-level recruitment of neurons for the processing of complex stimuli. The Inhibitory neurons are known to decorrelate network, enhance or restrict network plasticity (Tetzlaff et al., 2012; King et al., 2013). Thus, we hypothesize the role of inhibitory interneurons to understand the mechanism underlying differential adaptive coding of sound sequences.

The local Inhibitory interaction controls the gain modulation. Previous studies have shown divisive and subtractive modulations by optogenetically manipulating PV and SOM neurons, respectively (Seybold et al., 2015). Furthermore, PV and SOM neurons’ role during context-dependent gain modulation, such as in adaptation, has been actively studied (Natan et al., 2015b, 2017). Such studies still focus only on the local, short-term adaptation effects of PV and SOM on adapted and non-adapted neurons’ responses. We present an investigation, where we have focused on the effects of these inhibitory subtypes on the long-timescale adaptation and how statistical patterns within the stimuli affect their inhibitory control over the responses. Our results also show that such excitatory-inhibition connection takes longer than their excitatory counterparts to exert their inhibitory control on the responses (Fig. 6A). Our study observed that differential Inhibitory influence emerges over time, where EX-PV connections behave differentially to Periodic and Random sequences compared to EX-EX and EX-SOM connections, showing a denser functional connectivity for the periodic stimuli, unlike the EX-EX and EX-SOM connectivity, which had a denser functional connectivity for the random stimuli (Fig. 6A). In other words, as we know that inhibition is always preceded by excitation, owing to the recurrent and columnar architecture of the cortex; similarly, the adaptation properties of these inhibitory controls take place after the excitatory rebalancing of the responses. This could point out to a possible excitatory-inhibitory rebalancing act, for which the inhibitory neurons take longer than the excitatory ones. Our results suggest that EX-PV and EX-SOM connectivity has a strong role in long scale context-specific modulation, as seen by a change in connectivity pattern emerging in the later iterations. Within the EX-PV connections, the functional connectivity was much more homogenous and non-specific as compared to the EX-SOM and the EX-EX connections. Altogether, these results paint a vivid picture of the connections properties of the neural circuit which hints that the encoding of an aperiodic and random stimuli sequence require a more specific and directed connectivity profile with EX-EX and EX-SOM connections carrying most of the computational load as compared to the encoding of periodic and regular sound sequences, which requires relatively less specific and homogeneous connectivity and mostly rely on EX-PV connections. Such modifications throw a light on the differences in the modulating effects that are undertaken by the PV and SOM neurons.

We have discovered possible caveats in understanding how the auditory cortex identifies and encodes periodic and random stimulus patterns. Performing targeted optogenetic manipulation of PV and SOM neuron in future studies can further elucidate the role we decipher in functional subnetworks in the ACX. Nevertheless, we are confident that these results are a strong motivation to explore how statistical patterns are differentiated and discriminated by the auditory cortex. Overall, we see that our results, fall in synergy with the present understanding of this topic and opens up possibilities of inhibitory neurons role in the coding of sound sequences over long time scales for future research.

## Acknowledgements

MM thanks CSIR for Fellowship, AM thanks MHRD for PMRF, SB thanks the IIT Kharagpur Sponsored Research and Industrial Consultancy, Challenge Grant SGIGC-2015/DMN, and MHRD Signals and Systems for Life Sciences SSLS/VTA, scheme funds. *This work was supported by the DBT/Wellcome Trust India Alliance Fellowship IA/I/11/2500270 awarded to SB*.

## Material and Methods

### In-vivo extracellular recordings

#### Surgical procedures

Adult C57BL6/J JAX mouse (P30-P40) were initially anaesthetized under deep anaesthesia (5% isoflurane) inside the induction chamber. After that, during the time of surgery, the level of anaesthesia was lowered down to 1.5-2.5%. The internal body temperature was monitored and continuously maintained at 37°C using the heating pad throughout the experiment. The left temporal portion of the skull was exposed by performing a central incision and retracting the scalp. Tissue clearance and sterilization of the exposed area were performed by applying 3% H_2_O_2_ and alcohol, respectively. A craniotomy was made over an estimated left auditory cortical area bounded by the temporal ridge, lamboid suture, and ventral and rostral squamosal suture. Recordings were confirmed from A1 based on the direction of the tonotopic gradient.

### Recording procedure

Recordings were performed using the 4×4 multielectrode array (Microprobes, 125micron interelectrode spacing) of 3-5 Megaohms (MicroProbes, USA) impedance. L2/3 responses were probed between 200-300 microns from the cortical surface using a micromanipulator (MP-285, Sutter Instrument Company, Novato, CA). Signals were acquired after passing through unity gain (1x) headstage, followed by a preamp (Plexon, HST16o25) with 1000x gain. The wideband signal (LFP, 0.7Hz to 6 kHz) and spike signal (150Hz to 8Khz) were parallelly acquired through National Instruments Data Acquisition Card (NI-PCI-6259) at 20 kHz sampling rate. Further, offline/online analysis of obtained signals was performed using custom-written codes in MATLAB (Mathworks Inc. Natick, MA).

### In-vivo 2-photon imaging

To prepare a mouse for in-vivo 2-P imaging, standard surgical procedures were performed. Surgeries were performed under 1.5-2% of isoflurane anaesthesia. The left temporalis was exposed by transecting the scalp. Further, the exposed area was cleared and sterilized using 3% H_2_O_2_ and alcohol. A 5mm diameter circle was marked over the estimated auditory area. A metal plate was fixated using dental acrylic after keeping the mark in the centre of the head plate. A cranial window was created by lifting a skull flap of 3mm diameter over the left auditory area. After that, a craniotomy was filled with 1.5% low-melting agarose. Immediately after that, a 3mm coverslip was imbedded over the craniotomy to avoid brain pulsation. Animals were thereafter transferred to a soundproof chamber equipped with 2-photon imaging microscopy. Throughout the experimentation, the animal temperature was monitored continuously and maintained at 37°C. During the imaging sessions, anaesthesia was reduced down to 0.5-0.75% of isoflurane.

### Viral Injection

Adult mice (>40 days old, male and female) were anesthetized with isoflurane, as performed in extracellular recordings. Further, mice were injected with dexamethasone (2 mg/kg body weight) intraperitoneally and placed on a stereotaxic frame. The animal’s body temperature was maintained at 37 °C throughout the surgical procedure using a heating pad. An incision was made to expose the skull. A craniotomy of 3mm diameter was performed to expose the ACX, based on landmarks (rhinal vein). Virus (AAV.Syn.GCaMP6s.WPRE.SV40) was loaded into a glass micropipette mounted on a Nanoject II attached to a micromanipulator and injected at a speed of 20 nL per min at desired placed of ACX in PV-tdTomato and SOM-tdTomato transgenic animals. The craniotomy was covered with a glass coverslip. Mice were left in their home cage to recover and virus to integrate and express for two weeks. Experiments were conducted 2-3 weeks days after the viral injection.

### Sound delivery

Sounds were generated through TDT RX6, attenuated using TDT attenuators (PA5), and delivered by TDT electrostatic speakers (ES1) driven by TDT drivers ED1. Sounds were presented at a distance of 10 cm to the right ear. The frequency response of the ES1 speaker measured with a microphone 4939 (Brüel & Kjær, Denmark) showed a typical flat (+/-7 dB) calibration curve from 4-60 kHz. In a 2-P imaging setup, sounds were generated through TDT RZ6 multifunctional processor controlled by custom-written codes in MATLAB (Mathworks Inc. Natick, MA). The properties of each sound are given below:

Noise: Each 50 ms of 6-48KHz white band noise from 50-90 dB SPL with an interstimulus gap of ∼2 s was presented 5 times. The modest intensity of sound to be delivered was selected based on the majority of the recorded neurons sound level function.

Tone: Each 50 ms of pure frequency tone with frequencies ranging from 6kHz to 48kHz in 0.25-octave steps at 60-80 dB SPL (depending on noise threshold) and an interstimulus gap of ∼2 s was presented 5 times. Response to these tones was used to measure the best frequency and the tuning width of the neurons.

Oddball paradigm N-f pair: A series of sound tokens (n=15) each 50ms long (inter-token gap of 250 ms) of the pattern SSS…SDS…SS, with S as Standard and D as Deviant occurring with the ∼10% probability (1 in 15 tokens), were used for oddball characterization. A total of 30 iterations of such oddball sequences were presented with an inter-iteration gap of 2-8s. S and D consisted of pure tone, and broadband noise (6-48kHz with equivalent sound level). Responses were always collected in pairs with S and D swapped. The oddball sequences could be presented in the Periodic order or random order depending upon the position of the deviant. In random condition, deviant position was presented in random order appearing at a position anywhere between 2-15 token across all repetitions. However, in a Periodic condition, deviant always occurred at 8th token across all repeats.

Oddball paradigm f1-f2 pair. The sound structure is similar to the oddball paradigm N-f pair, as described previously. However, in this paradigm, all sequences consisted of pure tones with S and D as either f1 or f2. The two frequencies evoking high responses were selected, and the difference between f1 and f2 varied from 0.25-1.25 octave apart.

## Data Analysis

### Spike sorting

Spike sorting was performed using custom-written codes in MATLAB (Mathworks Inc. Natick, MA). Potential spikes were isolated based on fluctuations exceeding 4 standard deviations from the baseline from 8-16 independent channels. Further, spike waveforms were clipped from the raw data and projected into the space defined by the first three principal components. Clusters representing single-unit responses were identified by isolation distance between the cluster centroid quantified by K-mean clustering. The quality of the spike waveform was determined by visual inspection.

### Rate calculation

Further analysis was performed on the firing rate calculated within the stimulus duration or multiple tokens as in the case of the oddball sequence paradigm. A single unit was considered responsive if a response to at least one of the 4 stimuli (f/N or f1/f2 as Standard or Deviant) was significantly different from the baseline (300ms preceding stimulus, 2-tailed t-test, p<0.05). The inclusion criteria for the single-unit was the presence of significant response to at least one of the case (Standard/Deviant in Random and Periodic condition). Always paired data (Random and Periodic sequence) was considered for further analysis.

### Moving average analysis-extracellular recordings

We measured the long scale of adaptation to the standards and deviants in random and Periodic oddball sequences over 30 repetitions. For analyzing the change in firing rate across 30 iterations in random and Periodic sequences, 21 mean firing rates were calculated by performing a moving average of firing rates across 10 repetitions. A mean firing rate of all the standard token preceding deviant was considered for evaluating the trend in mean firing rate of standards across all the trials. Mean response of 7 tokens preceding the deviant was taken in the Periodic deviant (8th position) case; however, the mean response of the number of tokens to be considered is variable (1-14) in random deviant (2^nd^-15^th^ position) condition. To account for such biases in random sequences, we performed a weighted mean average response by taking a product of the number of tokens before deviant and the mean firing rate of all the tokens before deviant. After that in each moving average window, mean firing rate over 10 repetitions was divided by the sum of all standard tokens preceding deviant occurring in 10 repetitions of a given moving window. Normalized firing rates were computed over 21 moving averages by dividing it with the mean firing rate of the first moving average of 1-10 repetitions.

A slope across 21 moving average firing rates was evaluated using ‘regress’ function in MATLAB, to see how well the firing rates change over the trials. We considered all the slopes by dividing into 3 groups (+1, 0 and -1) for proportion analysis. A higher positive slope (+1) means firing rates increase significantly (F-test, p<0.05) in the later iteration, whereas 0 and -1 slope indicates no significant change or decrease in firing rate in the later repetitions, respectively. To compare Periodic and random sequences, the slopes’ inclusion criteria were the presence of a significant slope (F-test, p<0.05) in either Random or the Periodic sequence.

### Best Frequency

The best frequency of the neuron is the frequency which evoked the highest response within the stimulus duration (50 ms) to the same sound level at which the oddball paradigm was presented.

### CSI Index

To quantify a neuron’s selectivity to the deviant we used the common selectivity index (CSI).\

Tone(f1)-tone(f2) case: CSI = [*R*_*D*_(*f*_*1*_)+*R*_*D*_(*f*_*2*_)-*R*_*S*_(*f*_*1*_)-*R*_*S*_(*f*_*2*_)] / [*R*_*D*_(*f*_*1*_)+*R*_*D*_(*f*_*2*_)+*R*_*S*_(*f*_*1*_)+*R*_*S*_(*f*_*2*_)]

Noise(N)-Tone(f) case: CSI= [*R*_*D*_(*f*)+*R*_*D*_(*N*)-*R*_*S*_(*f*)-*R*_*S*_(*N*)] / [*R*_*D*_(*f*)+*R*_*D*_(*N*)+*R*_*S*_(*f*)+*R*_*S*_(*N*)]

where *R*_*S*_(.) and *R*_*D*_(.) denote response to Standard(S) and Deviant(D) respectively.

### Noise correlation

Noise correlation was computed between a pair of neurons recorded simultaneously in both random and Periodic deviant sequence. First, we represented a response to an oddball paradigm of an i^th^ neuron binned 50ms at time t to the n^th^ iteration as r_i,n_(t). Noise correlation was calculated between the i^th^ and j^th^ neuron by calculating the Pearson correlation coefficient between two simultaneously recorded spike trains as 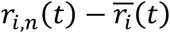 *and* 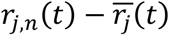, where 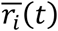is the mean response of i^th^ neuron over all the trials.

### LFP analysis

The raw electrical signals acquired at 20Khz sampling rate were baseline shifted to have a mean of baseline (200 ms preceding the stimulus) at 0. After baseline zeroing, the acquired signals were notch filtered (50Hz, Butterworth 8^th^ order, to remove AC supply line noise) and then bandpass filtered (between 1 Hz and 300 Hz, Butterworth 2^nd^ order). The processed signals were further resampled at 2kHz and stored for offline analysis. Response magnitude was quantified by considering RMS (Root mean square) of the potential fluctuations within the stimulus duration or multiple tokens as in case of oddball paradigm. Only those channels corresponding to which single-units that evoked significant response were considered for LFP analysis. Analysis performed on the LFP using RMS values was similar to spike analysis.

### Statistical tests-extracellular recordings

Chi-squared tests were used to compare the multiple ratios in the bar-plots showing the percentage of various slopes. Single value z-test was done to compare two individual ratios.

### Imaging analysis

#### Pre-processing

The fluorescence data collected from in-vivo 2-photon calcium imaging was sorted to selected to collect the mean fluorescence fluctuation within a 10mm circle centred around the cyton body of the neurons. The spatial mean fluorescence within that circle was considered to be the activity of that cell. This data was baseline zeroed by 1-1.5 seconds of spontaneous fluorescence data, to obtain the df/f values for every iteration separately.

### Best frequency calculation

The tonal frequency that elicited the maximum significant response with respect to the spontaneous was selected as the best frequency. The significance was calculated between the 4 frames preceding the onset and the mean activity within 500ms of the tone onset.

### Moving average analysis-2-P imaging

To find the standard response, the mean fluorescence from the onset frame to the frame just before the deviant was taken for each iteration. The deviant response was chosen as the deviant frame’s mean responses till just before the arrival of the next standard token. The standard and deviant responses from these 30 iterations was smoothened by applying an across iteration moving average window of 10 iterations with 1 iteration step, making a total of 21 data points for each cell and each stimulus. To find the slopes, we performed a linear fit on these data points, the cells with significant positive or negative slopes were designated a slope sign value of +1 and -1 respectively. All the other cells were designated a value of 0 slopes.

### Connectivity Analysis

To assert connectivity between two cells, we selected 30 iterations randomly from the original 30 iterations with replacement and computed the mean Pearson correlation between the two cells,this was done a total of 100 times and a 99% confidence interval was constructed from these values. A chance distribution was also constructed by shuffling the iteration between the two cells and performing the above steps. If the mean of the bootstrapped distribution was higher than the mean of the shuffled distribution by a 99% confidence interval, those two cells were considered to be connected (+1). In all the other cases, no connectivity was considered to be present (0). This method was used against the traditional noise correlation because we wanted to remove the uncertainty in the degree of connectivity between two given cells. This connectivity was found separately for the four different kinds of stimuli used in order to maintain their functional relevance.

### Gini Index

Having found the functional connectivity, we categorized the cells in 9 different categories, depending on the combination of slopes in response to the two different kinds of stimuli (3 slope types for noise standard multiplied by 3 slope types of tone standard). This was done separately for the connectivity pertaining to the Periodic stimuli and the random stimuli. In both the cases, if any pair was connected for at least one kind of stimuli (either noise or tone or both), we considered them connected. We generated a weighted graph with the nodes representing the 9 different categories of cells and the edges representing the number of connections between two given categories. Since we were more interested in studying the functional connectivity within the categories, we ignored the connections shared by the same category’s neurons (loops in a graph). We used the Gini index as a measure to quantify the sparsity of these graphs. To do that, we first generated a connectivity matrix ‘A’ from those graphs with A(i,j)=A(j,i)=number of connections shared between category i and category j neurons. Since we were ignoring all the loops, the diagonal elements were all zero. We then computed the sum of this matrix along the rows, thereby computing the total number of edges emerging from all the 9 nodes (note that A(i,j)=A(j,i), so the double-counting was included intentionally). These 9 values were arranged in ascending order and were divided by the total sum of all the 9 values (to make sure that the sum of all these values is 1). A Lorenz curve (curve showing the cumulative sum of all these values arranged in ascending order, starting at zero and ending at 1) was constructed, and the area under this curve was computed using trapezoidal approximation method (say B). Gini index is defined as G=1-2B. Intuitively, it can be seen that if the graph is connected sparsely, the cumulative distribution will turn out to be very narrow giving a lower value of B and hence a higher Gini index, thus if the neurons of this nine-category were densely connected with each other, we would see a lower value of Gini-index and if the inter-category connections were sparse, we would see a higher Gini-index. The raw graph, which consisted of all the connections between the nine categories, was taken and a total of 500 random replicates were constructed, where the edges were drawn at random with replacements. This was done to generate a 90% confidence interval of the Gini index to compare Periodic and random cases’ overall functional connectivity.

### Relative connectivity probability

We selected all the connected pairs from the population 500 times with replacement and found out the number of connections that fell in each category. Dividing it by the total number of connections gave us the probability of different categories of connections in each of the 500 trials and thus we generated a 99% confidence interval of the probabilities of all the connection categories. We next constructed a chance distribution by shuffling the cells and there generating pairs may not connect that in real. We calculated the relative connectivity probability by finding the difference between the means of these two distributions (µ_bootstrapped_-µ_chance_). Those connections whose mean differed from the chance mean by a 99% confidence interval (either higher or lower) were considered to be significant connections

### Statistical tests

All the bootstrap comparisons were made by seeing if the respective confidence intervals between the two groups being compared were overlapping or not. Chi-squared tests were used to compare the multiple ratios in the bar-plots showing the percentage of various slopes. Single value z-test was done to compare two individual ratios.

## Extended data Figures and Tables

**Figure E1-1.**
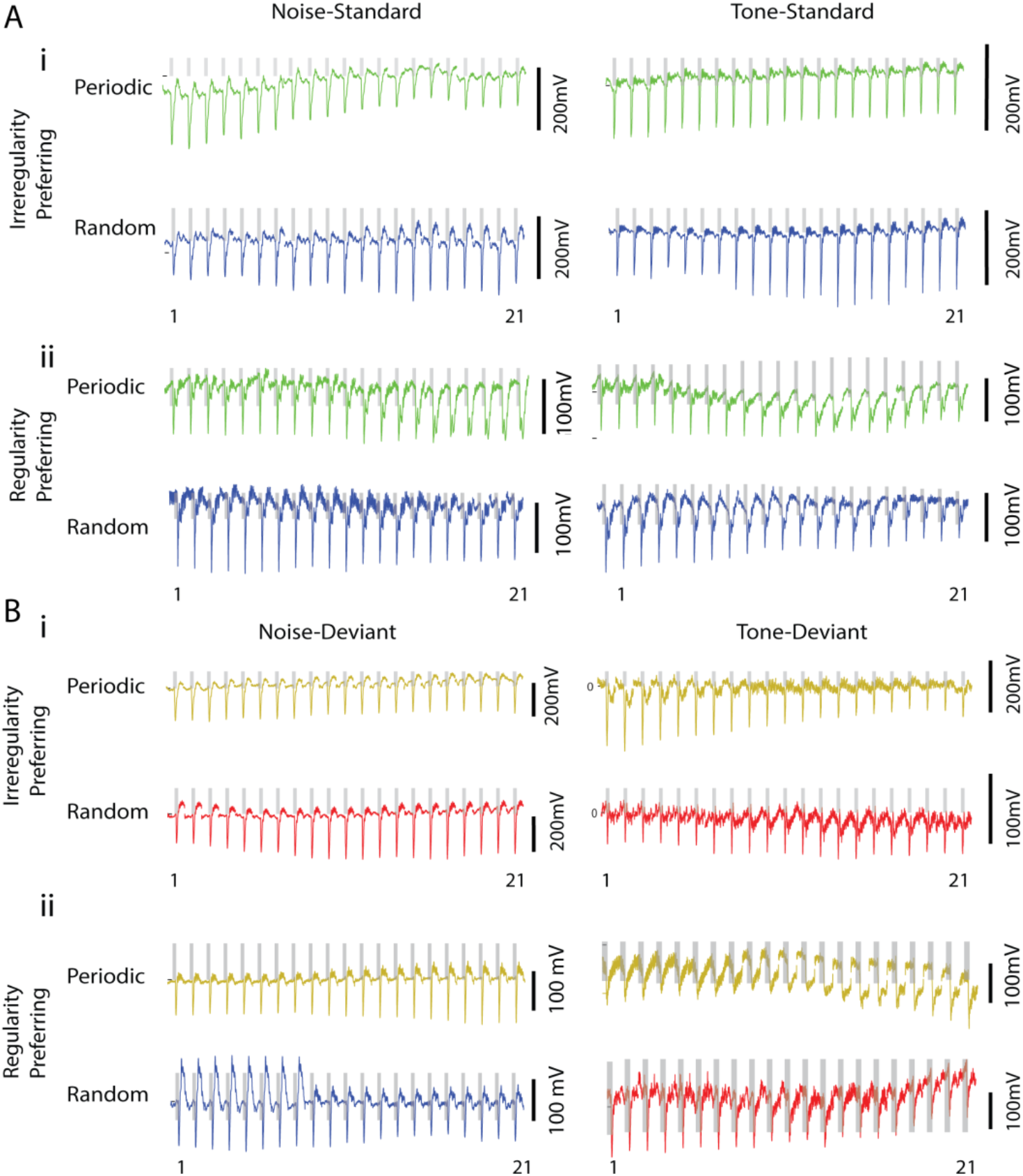
LFP response showing differential response pattern to periodic and random condition. A) Moving average LFP obtained from a single recording site to the Noise (Left) or tone (Right) as standards in Periodic (Green, Above) or Random (Blue, Below) condition. i) single-site recording example showing responses to standard tone and noise in Periodic oddball stimuli adapt over 21 moving averages, in contrast to an increase in response over later iteration in Random condition. ii) Another example showing opposite trend as shown in i), Responses to standard tone and noise in Periodic oddball stimuli increase over 21 moving averages, in contrast to an adaptation in response over later iteration in Random condition. Bi-ii) As shown in Ai-ii), single recording site examples illustrating over iteration LFP changes to deviant tone and noise in the periodic and random condition.

**Figure E2-1.**
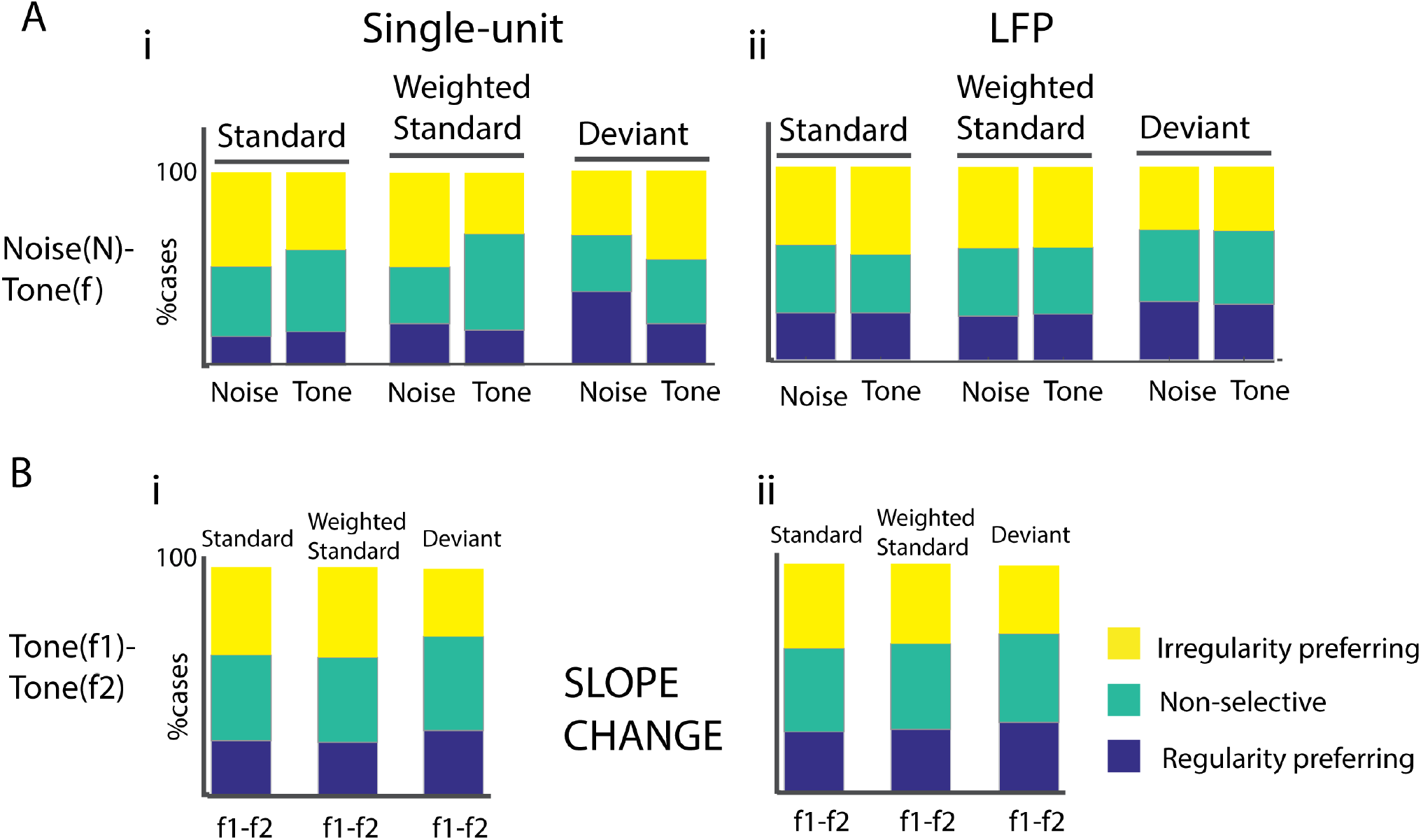
The proportion of Slope changes for tone and noise in Noise-tone and tone(f1)-tone(f2) oddball paradigm. A i) Change in slope (sign(slope_random_)-sign(slope_Periodic_) of single-unit across iterations moving average response for Noise and Tone as standards (Left), weighted standards (Middle) and deviant (Right) in Noise-tone oddball paradigm. ii) As shown in Ai, slope change of moving average LFP RMS for Noise and tone. B i-ii) As shown in Ai-ii, slope changes for f1-f2 as standard and deviant in tone (f1)-tone (f2) oddball paradigm.

**Table E2-1:** Population summary of data shown in Fig. S2Ai-ii, comparing Noise and Tone as standard and deviant in periodic and random case.

|  |  |
| --- | --- |
| Groups | Statistics: $\chi^2$ test for multiple proportion, followed by one sided z-propotionality test for individual slope values, 'T' denotes higher value in tone and 'N' denotes higher value in noise, followed by the p value of z-test. RP: regularity preferring, NS: non-specific, IP: irregularity preferring |
| Single unit: Noise (S) vs Tone (S) | $\chi^2=4.76$, $p=0.09$<br>RP: (N/T) 0.5; NS: (T) 0.0204; IP: (N) 0.0237 |
| Single unit: Noise (S) vs Tone (S), weighted | $\chi^2=14.07$, $p<10e-3$<br>RP: (N) 0.2314; NS: (T) 0.0001; IP: (N) 0.0013 |
| Single unit: Noise (D) vs Tone (D) | $\chi^2=14.41$, $p<10e-3$<br>RP: (N) 0.0001; NS: (T) 0.1005; IP: (T) 0.0099 |
| LFP: Noise (S) vs Tone (S) | $\chi^2=0.9$, $p=0.63$<br>RP: (N) 0.21; NS: (T) 0.22; IP: (N/T) 0.5 |
| LFP: Noise (S) vs Tone (S), weighted | $\chi^2=0.41$, $p=0.81$<br>RP: (N) 0.4; NS: (T) 0.26; IP: (N) 0.34 |
| LFP: Noise (D) vs Tone (D) | $\chi^2=0.68$, $p=0.7$<br>RP: (N) 0.33; NS: (T) 0.20; IP: (N) 0.34 |

**Figure E2-2.**
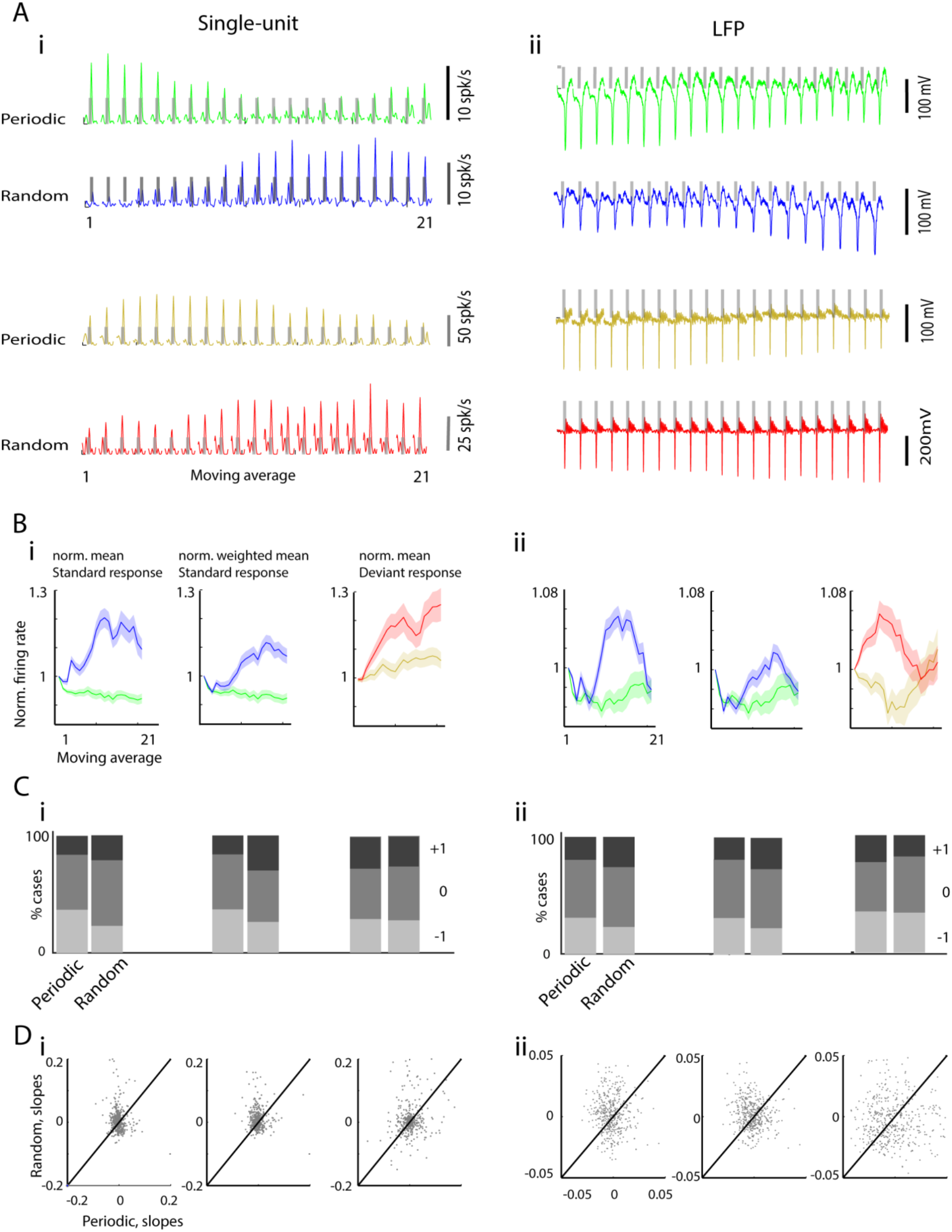
Response adaptation depends upon the structure of a sound sequence in tone (f1)-tone (f2) oddball paradigm. A i) Moving average Psth (Left) of a single Auditory cortex neuron and an LFP (Right) obtained in response to the tone as standards in Periodic (Green, Above) or Random (Blue, Below) condition. ii) Moving average Psth (Left) of a single Auditory cortex neuron and an LFP (Right) obtained in response to the tone as deviant in Periodic (Yellow, Above) or Random (Red, Below) condition. B i) The normalized mean population firing rate (Left); weighted mean population normalized firing rate to standards in Periodic and Random condition (Middle). (Right) Normalized mean population firing rate to a tone as deviant under two conditions. ii) Normalized mean population RMS obtained from LFP in response to tone as standard (Left and Middle) and deviant (Right) in f1-f2 oddball paradigm. C i) Proportion of -1, 0 and +1 slopes obtained from normalized and normalized weighted moving average firing rates to tone as standards (Left and Middle) and deviant (Right) in Random and Periodic condition. C ii) Proportion of -1, 0 and +1 slopes obtained from LFP RMS, as shown in C i). D i) The slopes obtained from moving average of firing rates in response to tone as standards (Left and Middle) and deviant (Right) under two conditions. D ii) As shown in D i), slopes obtained by calculating RMS from LFP.

**Table E2-2.**
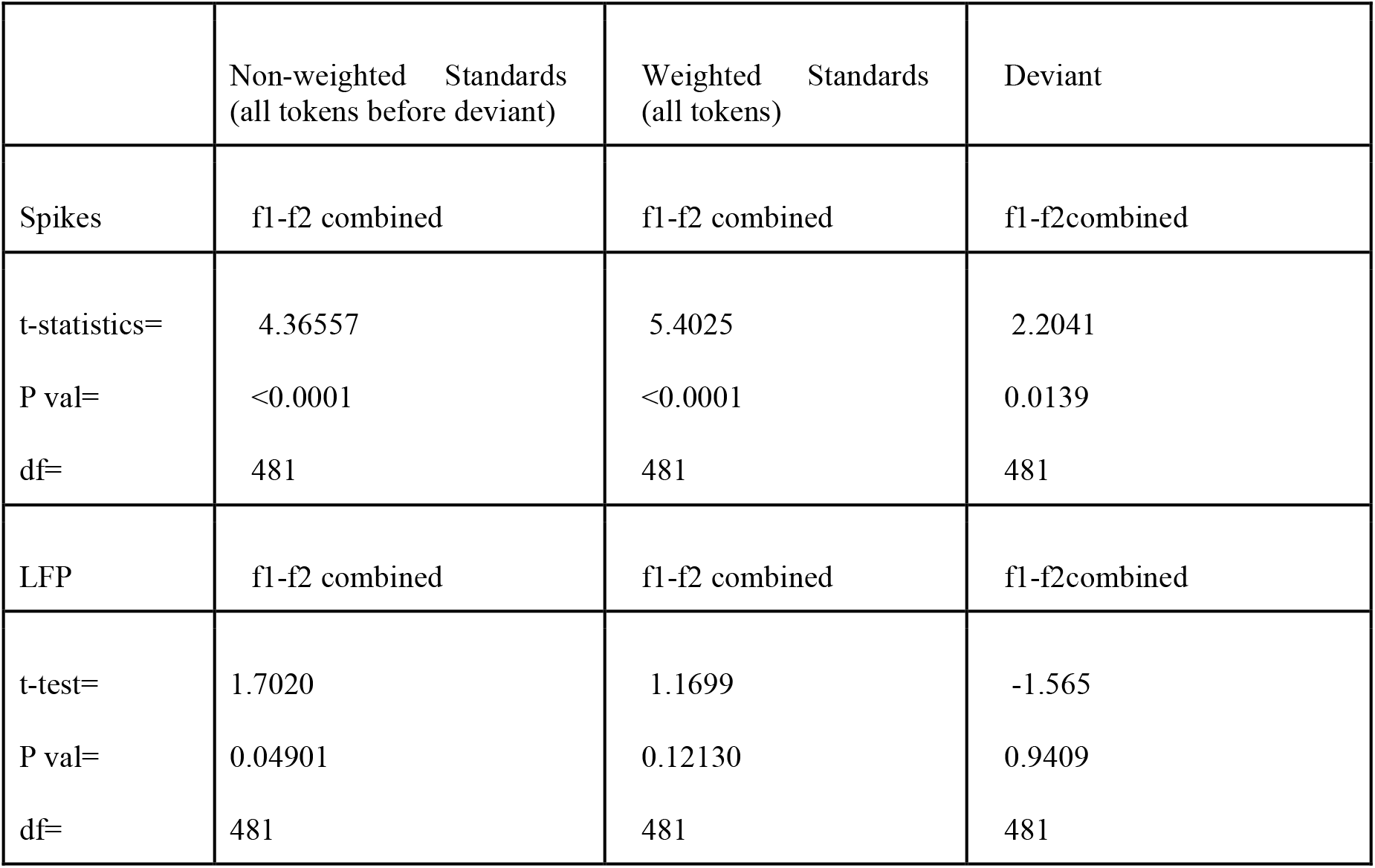
Population summary of data shown in Fig S1: Comparison of slope to f1-f2 combined in Periodic and Random sound sequence using tone(f1)-tone(f2) oddball paradigm.

|  | Non-weighted Standards<br>(all tokens before deviant) | Weighted Standards<br>(all tokens) | Deviant |
| --- | --- | --- | --- |
| Spikes | f1-f2 combined | f1-f2 combined | f1-f2combined |
| t-statistics= | 4.36557 | 5.4025 | 2.2041 |
| P val= | <0.0001 | <0.0001 | 0.0139 |
| df= | 481 | 481 | 481 |
| LFP | f1-f2 combined | f1-f2 combined | f1-f2combined |
| t-test= | 1.7020 | 1.1699 | -1.565 |
| P val= | 0.04901 | 0.12130 | 0.9409 |
| df= | 481 | 481 | 481 |

**Figure E2-3.**
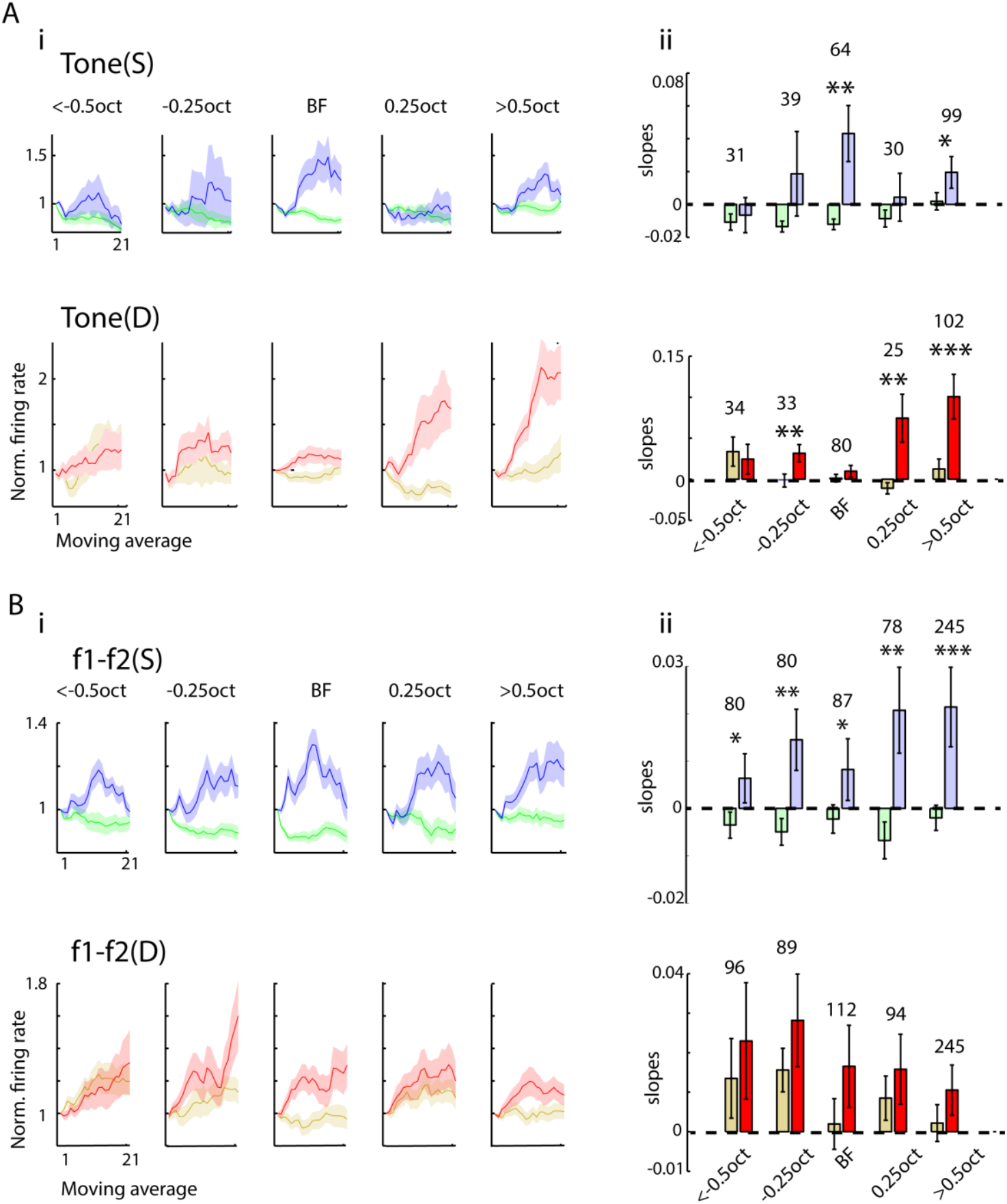
Moving average response as a function of distance from best frequency. A i) Population mean moving average response to Tone as Standard (Above) and deviant (Below) in Noise-tone oddball paradigm under two conditions, Periodic and Random at a different distance (octaves) from Best frequency (BF). ii) Population mean slope obtained as a function of distance from BF. B i-ii) As shown in (Ai-ii), Population mean moving average (Bi) and slopes (Bii) obtained in response to tone (f1-f2) as standard (Above) and deviant (Below) in tone(f1)-tone(f2) oddball paradigm. The number above paired-bars (Random-periodic mean slope comparison) corresponds to the degree of freedom. Colour scheme corresponds to the same as shown in Figure 1A. one-sided unpaired t-test, *p<0.05, **p<0.01, ***p<0.01.

**Figure E2-4.**
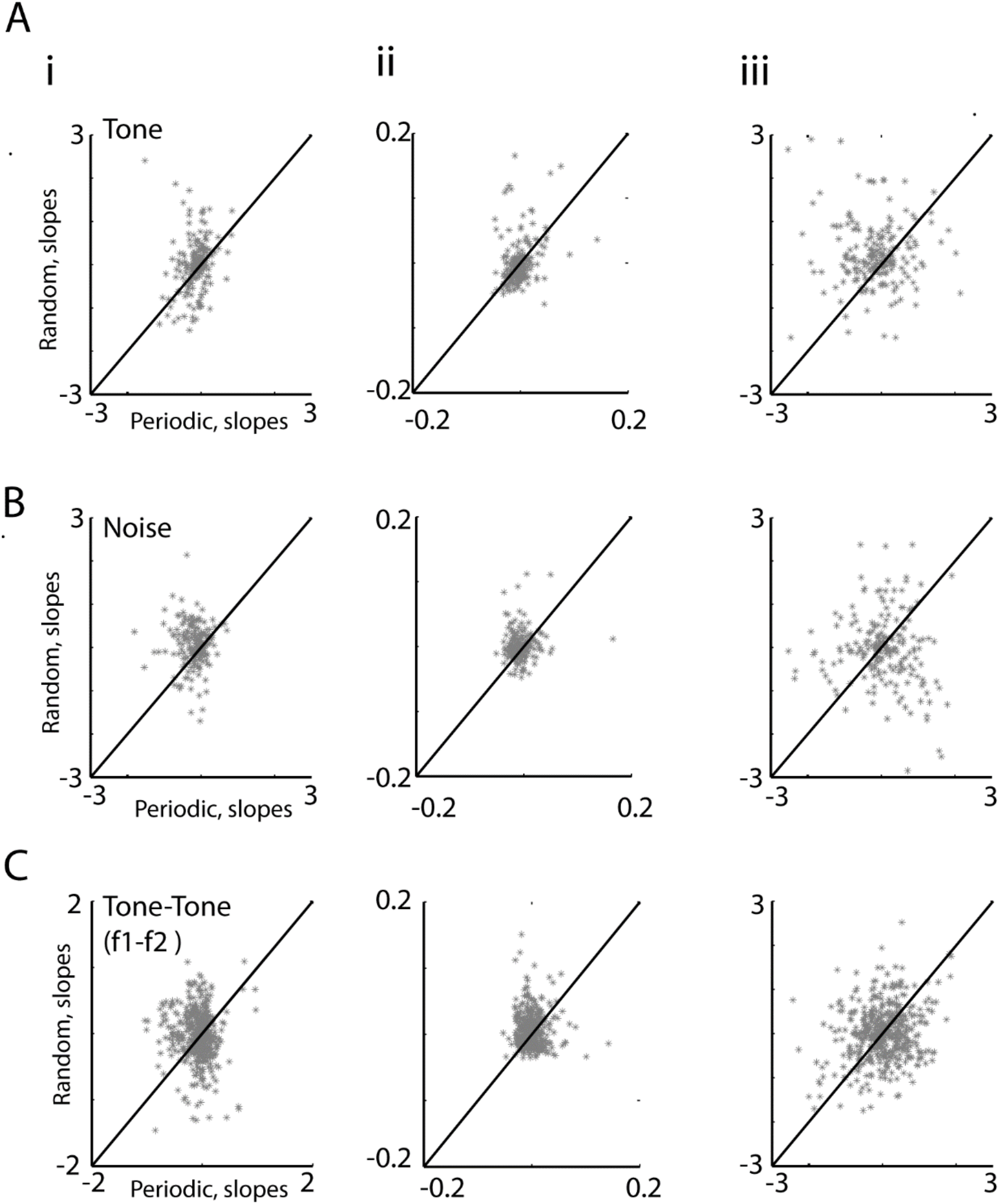
Differential moving average responses (slopes) to Standard and Deviant in Random condition and Periodic condition. i) Population slopes obtained from absolute firing rates (not normalized with first moving window) of single-unit moving average response to standard Tones (Ai), Noise (Bi) and tones(f1-f2 combined, Ci) in Periodic and Random condition. ii) Population slopes obtained from normalized firing rate of single-unit moving average response to all standard Tones (Aii), Noise (Bii) and tones(f1-f2 combined, Cii) in Periodic and Random condition. iii) Same as shown in (A-Ci), slopes obtained in response to deviant under two conditions presented.

**Table E2-4.** Population summary of Fig. S4: Comparison of slopes between Periodic and Random condition in different cases (Tone(f)-Noise (N) and Tone(f1)-Tone(f2)).

|  | The Non-normalized moving average response to only standards before dev token |  |  | Normalized moving average response: considering all standard tokens |  |  | The Non-normalized moving average response to deviant |  |  |
| --- | --- | --- | --- | --- | --- | --- | --- | --- | --- |
| Single -unit | f | N | f1-f2 | f | N | f1-f2 | f | N | f1-f2 |
| t-statistic= | 2.881 | 5.562 | 2.856 | 1.025 | 2.612 | 3.928 | 5.183 | -1.160 | 0.097 |
| P val= | <0.005 | <0.0001 | 0.002 | 0.153 | <0.005 | <0.0001 | <0.0001 | 0.876 | 0.538 |
| df= | 204 | 204 | 481 | 204 | 204 | 481 | 204 | 204 | 481 |

**Figure E2-5.**
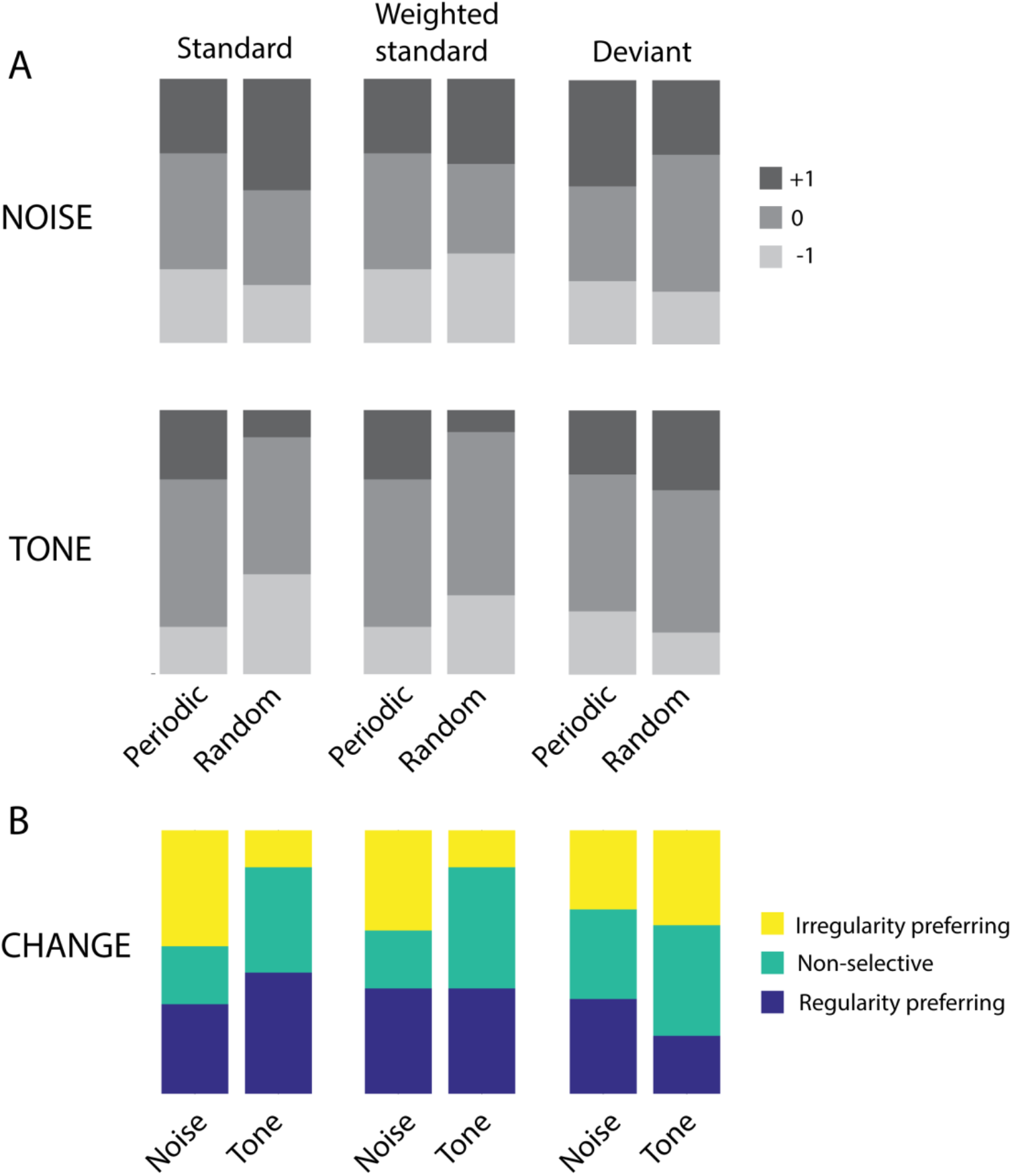
Slope distributions for single unit recordings for Noise-Tone stimuli in the thalamus (n=4 mice, 50 cases). A. Slope distribution for Tone and Noise stimuli as standard (left), weighted standard (Middle) and deviant (right). B. Change of slope distributions (sign(slope_random_)-sign(slope_Periodic_) for the two stimuli.

**Table E2-5a:** Population summary for data shown in Fig S5: Comparison of proportions of slopes in different cases.

|  |  |
| --- | --- |
| Groups | $\chi^2$ test for multiple proportion followed by one-sided z-proportionality test for individual slope values. The values in brackets denote a higher value in that category. 'P' denotes periodic; 'R' denotes random; 'N' denotes noise; 'T' denotes tone. RP:: regularity preferring, NS: non-specific, IP: irregularity preferring |
| Periodic standard (Noise) vs Random standard (Noise) | $\chi^2$: 2.16, p=0.34<br>-1: (P) 0.2442; 0: (P) 0.2071; +1: (R) 0.0711 |
| Periodic weighted standard (Noise) vs Random weighted standard (Noise) | $\chi^2$: 1.065, p=0.587<br>-1: (R) 0.258; 0: (P) 0.1527; +1: (R) 0.3313 |
| Periodic deviant (Noise) vs Random deviant (Noise) | $\chi^2$: 2.695, p=0.259<br>-1: (P) 0.3146; 0: (R) 0.0535; +1: (P) 0.1026 |
| Periodic standard (Tone) vs Random standard (Tone) | $\chi^2$: 7.2011, p=0.0273<br>-1: (R) 0.0130; 0: (P) 0.3441; +1: (P) 0.0187 |
| Periodic weighted standard (Tone) vs Random weighted standard (Tone) | $\chi^2$: 6.4172, p=0.0404<br>-1: (R) 0.08; 0: (R) 0.2709; +1: (P) 0.0083 |
| Periodic deviant (Tone) vs Random deviant (Tone) | $\chi^2$: 1.15; p=0.5621<br>-1: (P) 0.1587; 0: (R) 0.4206; +1: (R) 0.2496 |
| Change: Noise (Standard) vs Tone (Standard) | $\chi^2$: 11.27; p=0.0036<br>RP: (T) 0.11; NS: (T) 0.0258; IP: (N) 0.0005 |
| Change: Noise (Weighted standard) vs Tone (Weighted standard) | $\chi^2$: 9.77; p=0.0075<br>RP: (N/T) 0.5; NS: (T) 0.0057; IP: (N) 0.0031 |
| Change: Noise (Deviant) vs Tone (Deviant) | $\chi^2$: 2.38; p=0.3037<br>RP: (N) 0.0615; NS: (T) 0.2049; IP: (T) 0.2617 |

**Table E2-5b:** Population summary for data shown in Fig S5: Comparison of proportions of slopes and change of slope between Auditory cortex and Thalamus in different cases.

|  |  |
| --- | --- |
| Groups | $\chi^2$ test for multiple proportion followed by one-sided z-proportionality test for individual slope values (-1, 0 and +1). The values in brackets denote a higher value in that category. 'ACX' denotes Auditory cortex; 'Thal' denotes Thalamus. |
| Noise standard periodic | $\chi^2=14.94$, $p=10e-4$<br>-1(ACX): 0.003; 0(Thal): 0.0856; +1(Thal): 0.0013 |
| Noise standard random | $\chi^2=4.93$, $p=0.085$<br>-1(ACX): 0.4734; 0(ACX): 0.0308; +1(Thal): 0.0177 |
| Noise weighted periodic | $\chi^2=14.94$, $p=10e-4$<br>-1(ACX): 0.003; 0(Thal): 0.0856; +1(Thal): 0.0013 |
| Noise weighted random | $\chi^2=2.73$, $p=0.255$<br>-1(ACX): 0.3525; 0(ACX): 0.0648; +1(Thal): 0.0911 |
| Noise deviant periodic | $\chi^2=1.94$, $p=0.3776$<br>-1(ACX): 0.2936; 0(ACX): 0.2034; +1(Thal): 0.0815 |
| Noise deviant random | $\chi^2=3.09$, $p=0.21$<br>-1(ACX): 0.0399; 0(Thal): 0.1236; +1(Thal): 0.2988 |
| Tone standard periodic | $\chi^2=13.66$, $p=0.0011$<br>-1(ACX): 0.0002; 0(Thal): 0.0318; +1(Thal): 0.0127 |
| Tone standard random | $\chi^2=6.65$, $p=0.036$<br>-1(Thal): 0.13; 0(Thal): 0.1236; +1(ACX): 0.0050 |
| Tone weighted periodic | $\chi^2=13.66$, $p=0.0011$<br>-1(ACX): 0.0002; 0(Thal): 0.0318; +1(Thal): 0.0127 |
| Tone weighted random | $\chi^2=6.85$, $p=0.0325$<br>-1(ACX): 0.2087; 0(Thal): 0.0091; +1(ACX): 0.0197 |
| Tone deviant periodic | $\chi^2=3.24$, $p=0.198$<br>-1(ACX): 0.0462; 0(Thal): 0.0618; +1(Thal): 0.4651 |
| Tone deviant random | $\chi^2=2.68$, $p=0.2613$<br>-1(ACX): 0.3667; 0(Thal): 0.0546; +1(ACX): 0.0858 |
|  | RP: regularity preferring, NS: non-specific, IP: irregularity preferring |
| Change: Noise unweighted | $\chi^2=10.19$, $p=0.0061$<br>RP(Thal): 0.0011; NS(ACX): 0.029; IP(ACX): 0.272 |
| Change: Tone unweighted | $\chi^2=21.89$, $p=10e-5$<br>RP(Thal):$10e-6$; NS(ACX): 0.377; IP(ACX): 0.0003 |
| Change: Noise weighted | $\chi^2=6.9$, $p=0.0317$<br>RP(Thal):0.0043; NS(ACX): 0.152; IP(ACX): 0.085 |
| Change: Tone weighted | $\chi^2=12.72$, $p=0.0017$<br>RP(Thal):0.0006;NS(ACX):0.3169;IP(ACX):0.0063 |
| Change: Noise deviant | $\chi^2=0.4485$, $p=0.799$<br>RP(ACX):0.4189;NS(Thal):0.2567;IP(ACX):0.3340 |
| Change: Tone deviant | $\chi^2=1.8$, $p=0.4$<br>RP(Thal):0.4368;NS(Thal):0.1201;IP(ACX):0.1041 |

**Figure E3-1.**
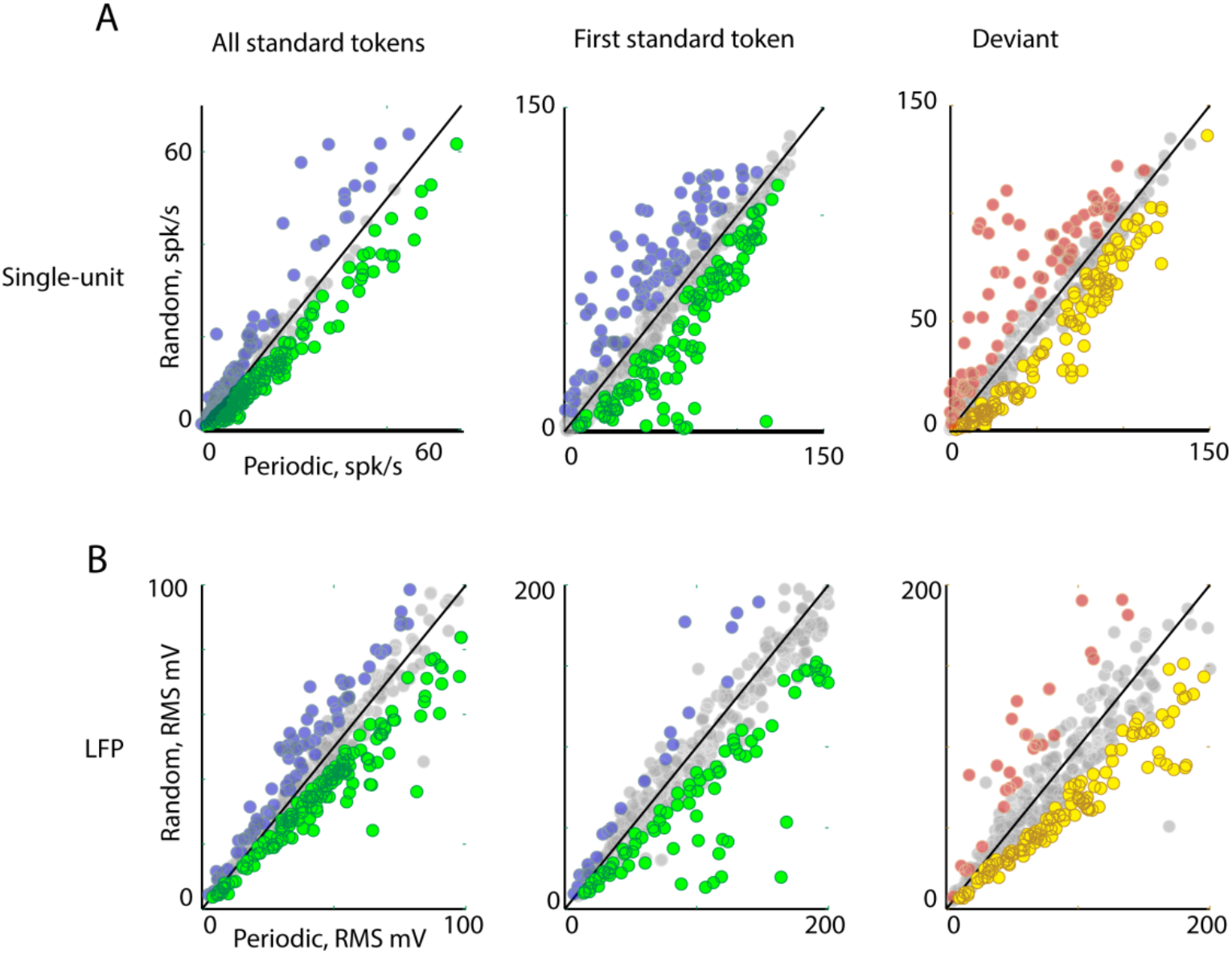
ACX L2/3 Single-unit and LFP responses to the periodic and random sequence. A) Top row: Population mean firing rate of cases to all f1-f2 standard tokens in Periodic and random condition (Left). Population mean firing rate of cases to first f1-f2 standard token in Periodic and random condition (Middle). Firing rate to f1-f2 as deviant under two conditions. B) Bottom row: Population mean RMS f1-f2 as standard (Left and Middle) and deviant (Right) in Periodic and Random condition.

**Table E3-1.** Population Summary: Responses to tone(f1-f2) as standard and deviant in tone-tone oddball paradigm under two conditions, Periodic and Random.

| LFP (RMS) | Standard all tokens (f1-f2) | Standard first token (f1-f2) | Deviant token (f1-f2) |
| --- | --- | --- | --- |
| Number of cases Periodic>Random | 303/482 | 317/482 | 303/482 |
| Fraction | 57% | 66% | 63% |
| T-statistics | 3.3776 | 7.4396 | 5.5887 |
| df | 481 | 481 | 481 |
| P val | 0.0003 | <0.0001 | <0.0001 |
| L2/3 Single unit | Standard all tokens (f1-f2) | Standard first token (f1-f2) | Deviant token (f1-f2) |
| Number of cases<br>Periodic>Random | 304/482 | 274/482 | 280/482 |
| Fraction | 64% | 57% | 58% |
| T-statistics | 4.6895 | 2.5585 | 2.3922 |
| df | 481 | 481 | 481 |
| P val | <0.0001 | 0.0054 | 0.0085 |

**Figure S4-1.**
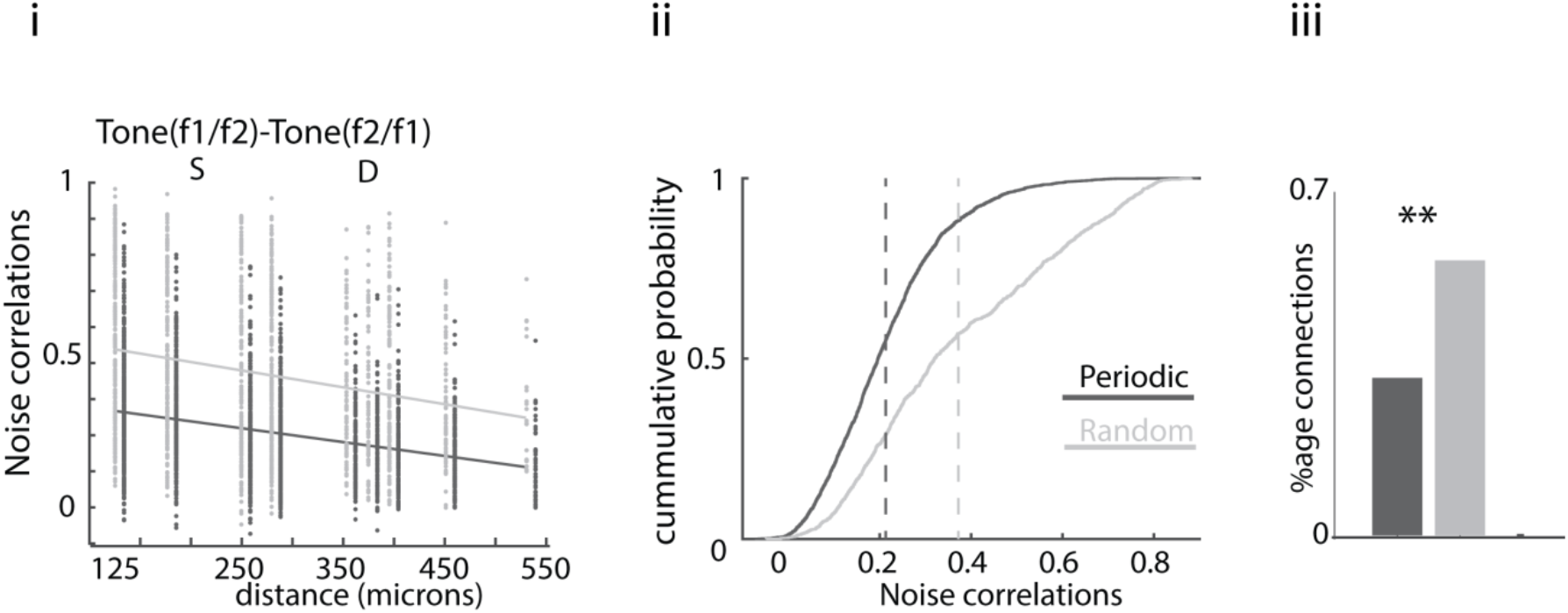
Random and Periodic condition differentially modulates Noise correlation and network connectivity. As shown in Main Text, Figure 4A i-ii, the noise correlation as a function of distance and distributions of obtained correlation coefficient to f1-f2 oddball stimulus in Periodic and Random condition. iii), Proportion of connection obtained in Periodic and Random condition using Granger causal analysis.

**Figure E5-1.**
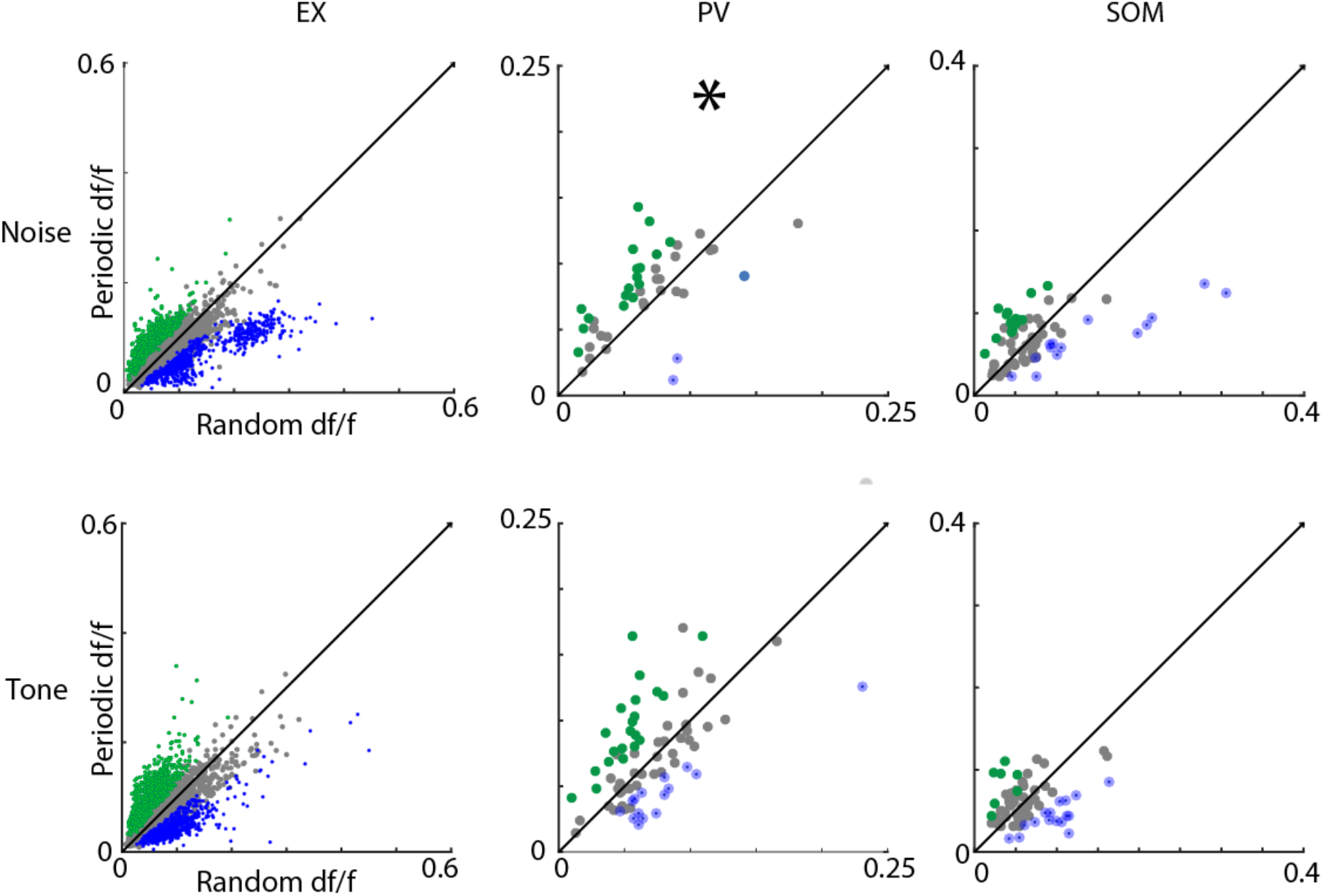
Scatter plot showing the mean df/f values for periodic and random stimuli in Noise and Tone as standard for the three neuron types. Table 5.1 below shows the statistics for every case. The coloured filled circles represent cases where a significant difference (paired t-test, p<0.05) of sound-evoked activity was observed between Random and Periodic condition. Color scheme corresponds to the same as used in Fig. 3-1.

**Table E5-1.** **Population Summary showing a comparison of mean df/f values between Periodic and Random conditions**.

| Cell type | Stimuli | Stats: One-sided paired t-test |
| --- | --- | --- |
| Ex | Noise | $(df/f)_{\text{random}} > (df/f)_{\text{periodic}}$; $t(6175)=1.1003$; $p=0.1356$ |
| Pv | Noise | $(df/f)_{\text{random}} < (df/f)_{\text{periodic}}$; $t(77)=1.9672$; $p=0.0264$ |
| Som | Noise | $(df/f)_{\text{random}} < (df/f)_{\text{periodic}}$; $t(101)=0.3598$; $p=0.3599$ |
| Ex | Tone | $(df/f)_{\text{random}} < (df/f)_{\text{periodic}}$; $t(6175)=1.2761$; $p=0.1010$ |
| Pv | Tone | $(df/f)_{\text{random}} < (df/f)_{\text{periodic}}$; $t(77)=1.1456$; $p=0.1277$ |
| Som | Tone | $(df/f)_{\text{random}} < (df/f)_{\text{periodic}}$; $t(101)=0.0731$; $p=0.4709$ |

**Figure E6-1.**
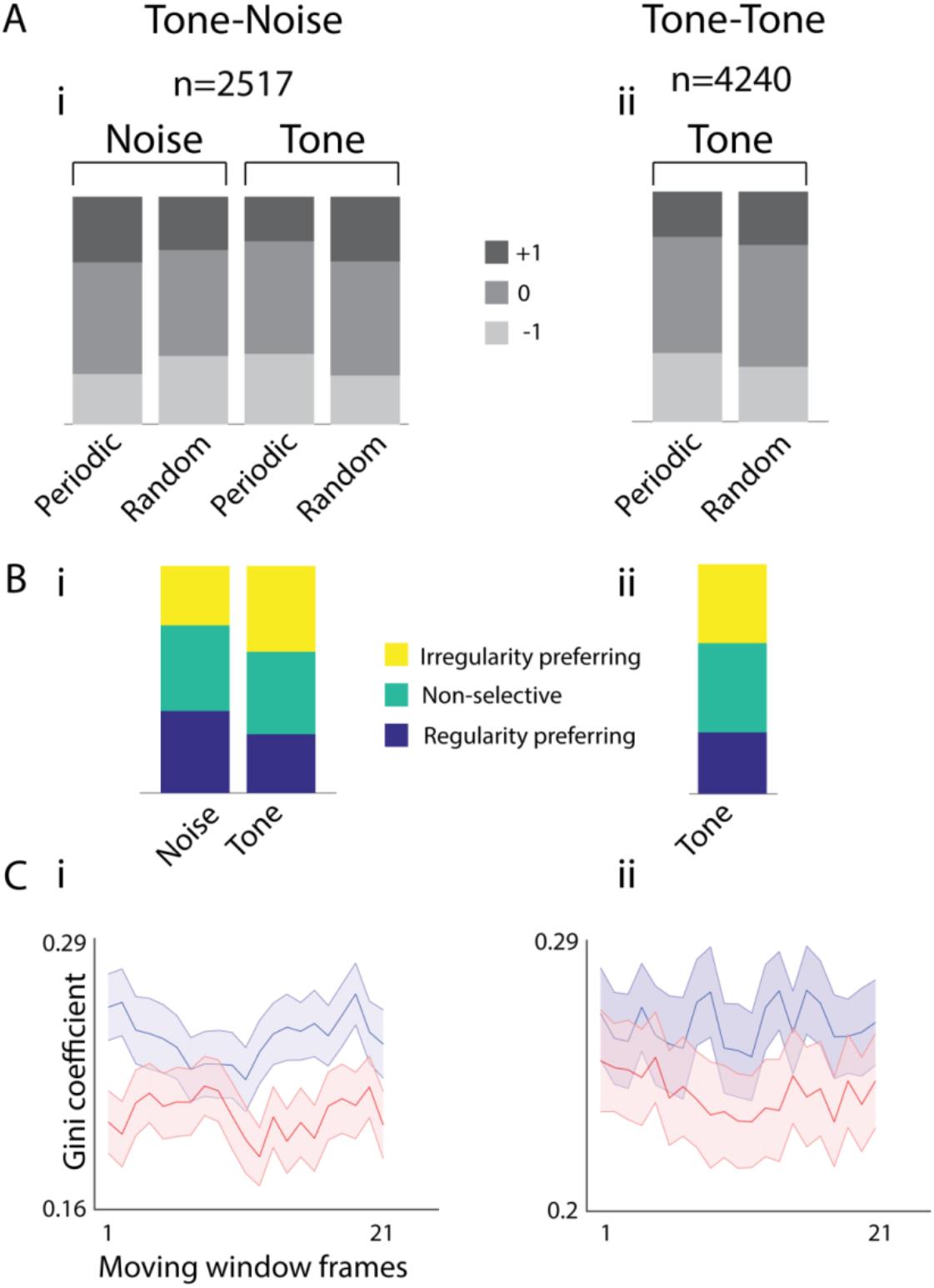
Slope distribution and Gini-coefficient for GCamp-Thy-1 animals. A) Slope distribution for i) Noise-Tone and ii) Tone(f1)-Tone(f2). The number above the bars indicate the number of neurons. B) Distribution of change of slopes for i) Noise-Tone and ii) Tone(f1)-Tone(f2). C) Gini-coefficient plots indicating functional connectivity in a moving average of 10 frames for periodic, fixed deviant (blue) and random deviant (red) case for i) Noise-Tone and ii) Tone(f1)-Tone(f2).

